# Active site mutations of F_420_-dependent alkene reductases reverse stereoselectivity

**DOI:** 10.1101/2023.02.21.529347

**Authors:** Suk Woo Kang, James Antoney, David W. Lupton, Robert Speight, Colin Scott, Colin J. Jackson

## Abstract

Ene-reductases from the Flavin/Deazaflavin Oxidoreductase (FDOR) family have potential value in biocatalysis as they typically exhibit complementary stereoselectivity to the widely utilized Old Yellow Enzyme (OYE) family, yet they are comparatively poorly understood at a mechanistic level. Here, we use a rational design approach to generate a library of 46 active site mutants of two FDORs from *Mycobacterium smegmatis* and examine the effects on conversion and stereoselectivity against a panel of substrates. Analysis of the effects of these mutations on stereoselectivity across all substrates revealed that the catalytic mechanism is highly sensitive to the polarity of the immediate active site. A conserved active site tyrosine in these enzymes, which does not serve as the proton donor, strongly affects stereochemical outcomes with Cα- (but not Cβ-) substituted substrates. Notably, a Tyr-Met mutation at this position reversed the diastereomeric excess (*de*) with (*R*)-carvone from 85.3% to −17.3% (cis/trans). Additionally, this mutation significantly increases activity with (1*S*)- verbenone. Finally, we show that the altered stereoselectivity is not due to a “flipped” substrate binding mode in these mutants, but rather that the hydrogenation mode is altered to favor *syn* relative to *anti* addition. These results show that the FDORs are highly engineerable and that, despite their superficial similarity, the OYE and FDOR families differ in crucial mechanistic aspects.

## Introduction

New and improved biocatalysts continue to be developed to complement traditional chemical synthesis in the manufacturing of flavors, fragrances, and pharmaceuticals.^1,2^ An important chemical transformation is the reduction of C=C bonds, as this may introduce up to two stereogenic centers into a molecule, often with high levels of stereoselectivity. Traditional synthetic methods require flammable (and often pressurized) H_2_ gas as well as expensive transition metal catalysts for which natural reserves continue to dwindle.^2^ Enzymes offer a compelling alternative as they are inherently renewable, highly active, operate at ambient temperatures and pressures, and generate relatively benign waste. Asymmetric reduction of alkenes conjugated to electron-withdrawing groups is catalyzed by several families of enzymes, of which the Old Yellow Enzyme family (EC 1.6.99.1) have been most thoroughly explored and utilized.^3–5^ These enzymes transfer the equivalent of H_2_ from NADH or NADPH to the substrate *via* the noncovalently bound cofactor FMN. Other flavin-independent families of ene-reductases that transfer the equivalent of H_2_ from nicotinamides directly to the substrate have been investigated, however these show comparatively narrow substrate ranges and modest chemoselectivity.^6–8^

More recently, ene-reductases that utilize the deazaflavin cofactor F_420_ have been shown to reduce their substrates in a regio- and enantioselective manner that is often stereocomplementary to OYEs.^9–11^ These enzymes belong to the flavin/deazaflavin oxidoreductase (FDOR) family of split β-barrel proteins.^12^ While the cofactor F_420_ occurs only in a limited number of prokaryotes, elucidation of its biosynthesis and metabolic engineering has enabled its production in the industrial workhorse *Escherichia coli* (in which it is not natively produced) at high titers, creating opportunities for its use as a bio-orthogonal cofactor in biocatalysis.^13–16^

F_420_-dependent FDORs exhibit remarkable diversity of substrates upon which they act, ranging from physiological substrates such as quinones, thiopeptins and tetracyclines, to promiscuous activities against aflatoxins, triarylamine dyes and nitroimidazole prodrugs.^17–23^ Several earlier reports have shown that that promiscuous ene-reductase activity is widespread amongst these enzymes.^24^ We recently reported detailed characterization of the ene-reductase activity and substrate range of the FDOR enzymes from the -A and -B subgroups.^10, 11^ Among the enzymes we have characterized, the FDOR-A enzymes MSMEG_2027 and MSMEG_2850 from *Mycobacterium smegmatis* demonstrated the best combination of activity and broad substrate range.^10^ Using X-ray crystallography and computational substrate docking it was possible to identify the preferred binding mode of substrates in the active site, which when combined with the experimentally observed products suggested that reduction proceeds with net *anti* addition of H_2_ across the double bond, as is most commonly observed in the OYE family.^3, 25–28^

Although mutagenesis has been used to interrogate FDORs involved in specific physiologically relevant activities such as reduction of biliverdin and antitubercular nitroimidazole prodrugs,^18, 20^ there are few reports of engineering FDORs as biocatalysts for the reduction of alkenes or using mutagenesis to probe their structure-function relationships in a broad manner. Here, we use a rational design approach to examine the active site residues of MSMEG_2027 and MSMEG_2850 involved in substrate binding and probe the effect of 46 unique active site mutants on activity and stereoselectivity with a panel of 10 substrates (460 separate mutant-substrate activities). Analysis of these results has identified multiple variants that exhibit enhanced activity and selectivity with the various substrates. In addition, several variants that show a reversal in stereoselectivity have been identified.

## Results

### Rational design of mutants

We previously reported the crystal structures of MSMEG_2027 bound with F_420_ (PDB 6WTA) and MSMEG_2850 in the apo state (PBD 8D4W) from which we generated holoenzyme models and performed computational substrate docking.^10^ For our rational protein engineering approach, we selected (*R*)-(-)-carvone (1a) as a model substrate because (i) the bulky 5-substituent should strongly direct binding orientation; (ii) the opposite enantiomer (2a) is also available; (iii) it is a widely used substrate for studies of ene-reductases; and (iv) the wildtype enzymes showed moderate to high conversion and good but not complete stereoselectivity, and mutations that increase and/or invert selectivity could therefore be identified.

We opted to use a rational design approach to leverage the available structural information, as well as to reduce the number of variants that would need to be screened using low-throughput GC/MS methods. Our approach used a combination of molecular docking and molecular mechanics generalized born surface area (MM-GBSA) as implemented in the Schrodinger Biologics suite of programs. MM-GBSA has been used previously in engineering transaminases to accept bulky substrates through identification of mutations that increased the relative binding affinity of the substrate.^29^ Using the previously obtained docking poses of **1a** (Figure 1) we selected residues for mutation. A total of thirteen and fourteen residues of MSMEG_2027 and MSMEG_2850, respectively, were within 5 Å of the docked ligand. This included the SKGG motif that is highly conserved in the FDOR-A family, where the amide NH of G69 and the hydroxyl of S67 (numbering for MSMEG_2027) form an oxyanion hole where the substrate carbonyl (or equivalent) binds (Figure 1A).^10, 12^ We selected the S67A mutant of MSMEG_2027 to examine the contribution of the serine hydroxyl to catalysis but did not select further mutations in this region owing to strong conservation and the lack of involvement of sidechains in substrate binding. Of the remaining nine and ten residues of MSMEG_2027 and MSMEG_2850, respectively, we targeted those in loop regions: V30 and L31 in loop 1; M54 in loop 3; V65 in loop 5; Y120 and P122 in loop 10 (numbering for MSMEG_2027). Other selected residues were in α-helices: W11 and V12 in α1; Y126 in α4 (numbering for MSMEG_2027). The orientation of the side chains of these target residues is shown along with the structural alignment (Figure 1B).

**Figure 1.**
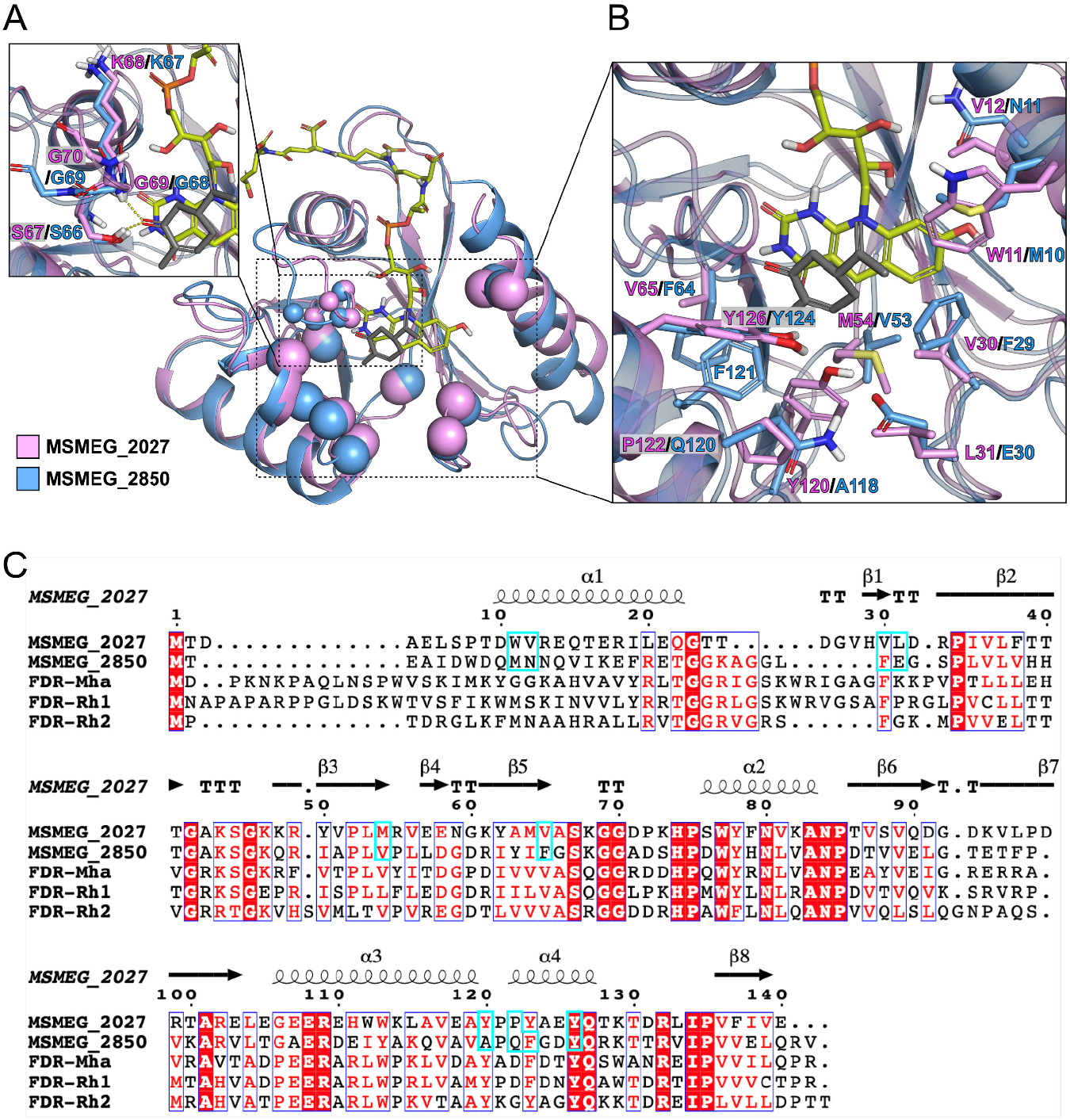
Choice of mutation sites for MSMEG_2027 (pink) and MSMEG_2850 (blue). The holo-enzyme structures complexed with F_420_ and (*R*)-(-)-carvone (**1a**, gray) were modelled based on crystal structures (PDB 6WTA and PDB 8D4W). Spheres represent Cα atoms of the residues located within 5 Å distance of **1a** in the active site. Cα atoms are depicted as smaller spheres in the SKGG motif. For clarity, only the F_420_-4 molecule of 6WTA and **1a** computationally docked into MSMEG_2027 are shown. **1a** and F_420_ appear in gray and yellow green, respectively. (A) The side chains of SKGG motif forming oxyanion hole with **1a** are shown as sticks. Hydrogen bonds are represented by yellow dashed lines. (B) The side chains of the mutational targets are shown as sticks. (C) Multiple sequence alignment of MSMEG_2027, MSMEG_2850, FDR-Mha, FDR-Rh1, and FDR-Rh2. Positions chosen for mutagenesis are indicated by cyan boxes.

Mathew *et al*., previously reported enantioselective ene-reduction by three other actinobacterial enzymes, FDR-Mha, FDR-Rh1, and FDR-Rh2 that also belong to the FDOR-A1 subgroup.^9^ We compared the active site residues using multiple sequence alignment and AlphaFold2 models^30, 31^ (Figures 1C and S1). While most residues in the active site are conserved across these FDOR-As, V30 and M54 are specific to MSMEG_2027 while F64 and A118 are specific to MSMEG_2850. Pairwise sequence identity ranged from 30.2 to 49.1% (Figure S2). These enzymes also contain conservative mutations in the SKGG motif that serves as the oxyanion hole (SQGG in FDR-Mha and FDR-Rh1, SRGG in FDR-Rh2). This position is less strongly conserved than the serine and second glycine.^12^

Each of the remaining target residues was mutated to the other 19 natural amino acids *in silico* using the Residue Scanning workflow in Schrodinger. Mutations that increased the predicted binding affinity in a catalytically productive binding mode (one in which the Cβ was within 3 Å of C5 of F_420_ following MM-GBSA refinement) were selected for experimental characterization. In the absence of a predicted improvement in binding affinity, favorable effects on protein stability and Prime energy (total system energy) were also taken into consideration.^32^ Altogether, 20 and 26 single mutations of MSMEG_2027 and MSMEG_2850, respectively, were selected for our mutant library, which is shown with computed parameters in Tables S1 and S2.

With the targeted library designed, constructs were generated, expressed, and purified. All variants gave high soluble expression and were purified to apparent homogeneity as judged by SDS–PAGE analysis (Figure S3). These were then assayed against **1a** in coupled enzyme assays using F_420_-dependent glucose-6-phosphate dehydrogenase to regenerate F_420_H_2_.

### Effect of mutations on activity with model substrate (*R*)-carvone (1a)

For MSMEG_2027, marginal increases in substrate conversion were observed with the V12N, V30F/W/Y, L31F/M/Y, V65L, and P122M/R variants, with V65L showing the highest increase in conversion (94.3% *cf*. 89.1% for the wildtype; Figure 2A and Table S3). Substantial reduction in substrate conversion was observed with the variants V30R, Y120Q and Y126M (34.8%, 19.3%, and 57%, respectively), while no conversion occurred with the M54Q and S67A variants. With MSMEG_2850, many variants showed higher conversion compared to the wildtype (84.2%), with V53I, A118F and Q120Y performing the best with 93.0%, 90.8% and 95% conversion, respectively (Figure 2A and Table S4). Substantial loss of activity was observed with the M10R, F29R, and F64N mutants (21.8%, 4.7%, and 7.6%, respectively) while no conversion was observed with the N11W, N11Y or F121Y mutants.

**Figure 2.**
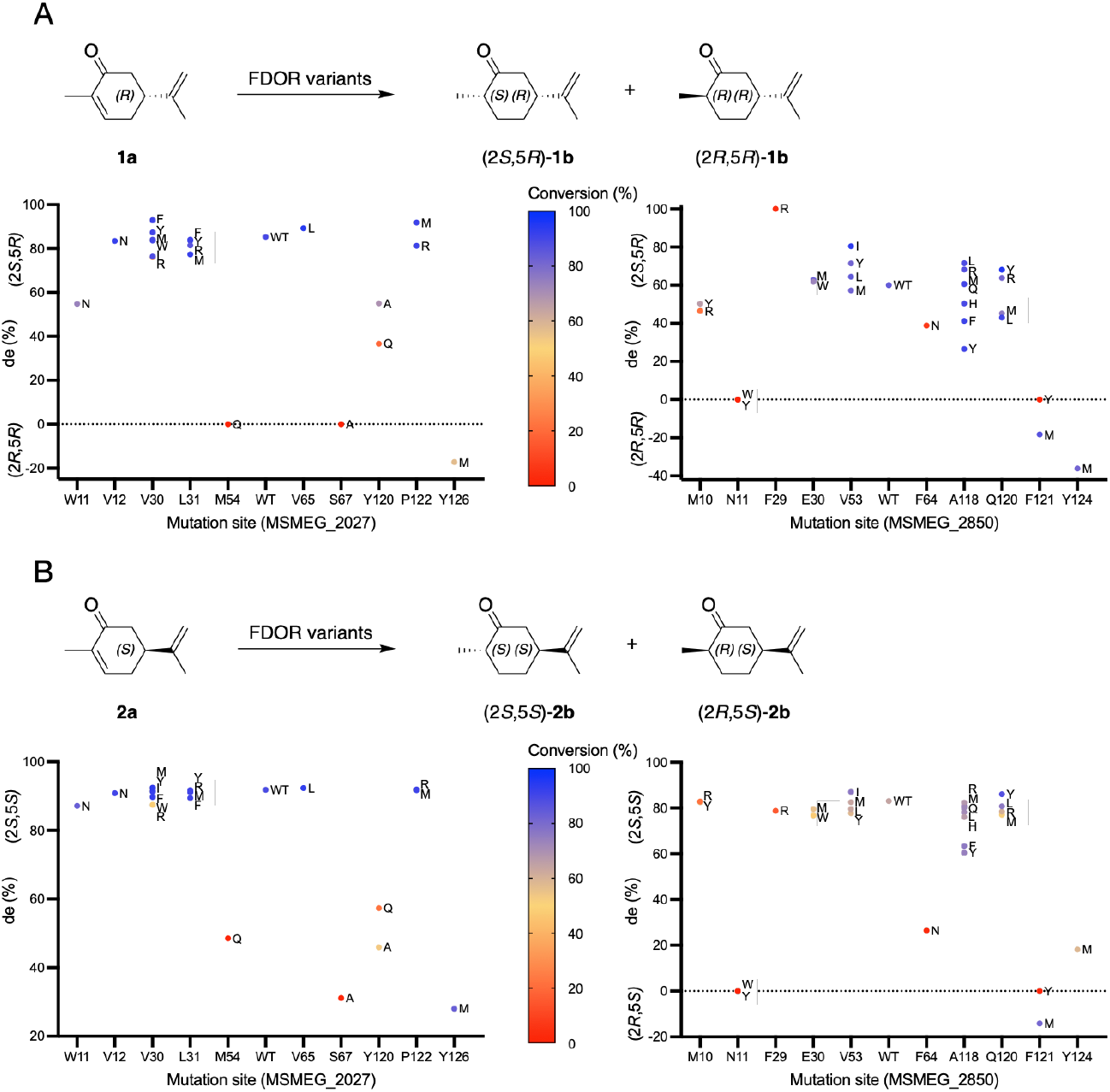
Performance of MSMEG_2027 and MSMEG_2850 variants in biotransformation of (A) (*R*)-carvone **1a** and (B) (*S*)-carvone **2a**. Mutants are grouped by residue number along the *x* axis and diastereomeric excess shown on the *y* axis. Conversion is indicated by fill color.

To our surprise, weak but reversed selectivity was observed with the variants MSMEG_2027-Y126M, MSMEG_2850-Y124M and MSMEG_2850-F121M (−17.3%, −36.2% and −18.3% *de*, respectively *cf*. 85.3% and 60.0% *de* for wildtype MSMEG_2027 and MSMEG_2850, respectively, Figure 2 and Tables S3 and S4). Y126 in MSMEG_2027 and Y124 in MSMEG_2850 are equivalent conserved positions (Figure 1) and the Tyr→Met substitution had similar effects in both enzymes. To our knowledge, this is the first report of FDORs mutants that exhibit altered stereoselectivity.

Multiple reports of stereocomplementary variants of OYE family ene-reductases have found that the altered selectivity can result from the substrates binding in an alternative “flipped” binding mode that results in hydride transfer to the opposite face of the double bond compared to the wildtype.^26, 27, 33, 34^ To investigate whether these mutations lead to “flipped” binding of **1a**, models of the Tyr→Met variants previously generated *in silico* were refined using MM-GBSA in the absence of the ligand. **1a** was then docked into these models using the induced fit protocol in Schrodinger. We found that the most energetically favorable pose of **1a** in these mutants had the same binding mode as the wildtype enzymes (Figure 3). No poses corresponding to the “flipped” mode of **1a** were observed in MSMEG_2027-Y126M, whereas no “flipped” poses were observed with MSMEG_2850-Y124M until the 9^th^ highest ranked pose. This pose had a docking score 1.1 kcal mol ^-1^ above that of the first ranked pose (−5.919 versus −7.005) This suggests that the altered selectivity did not result from a change in the binding mode.

**Figure 3.**
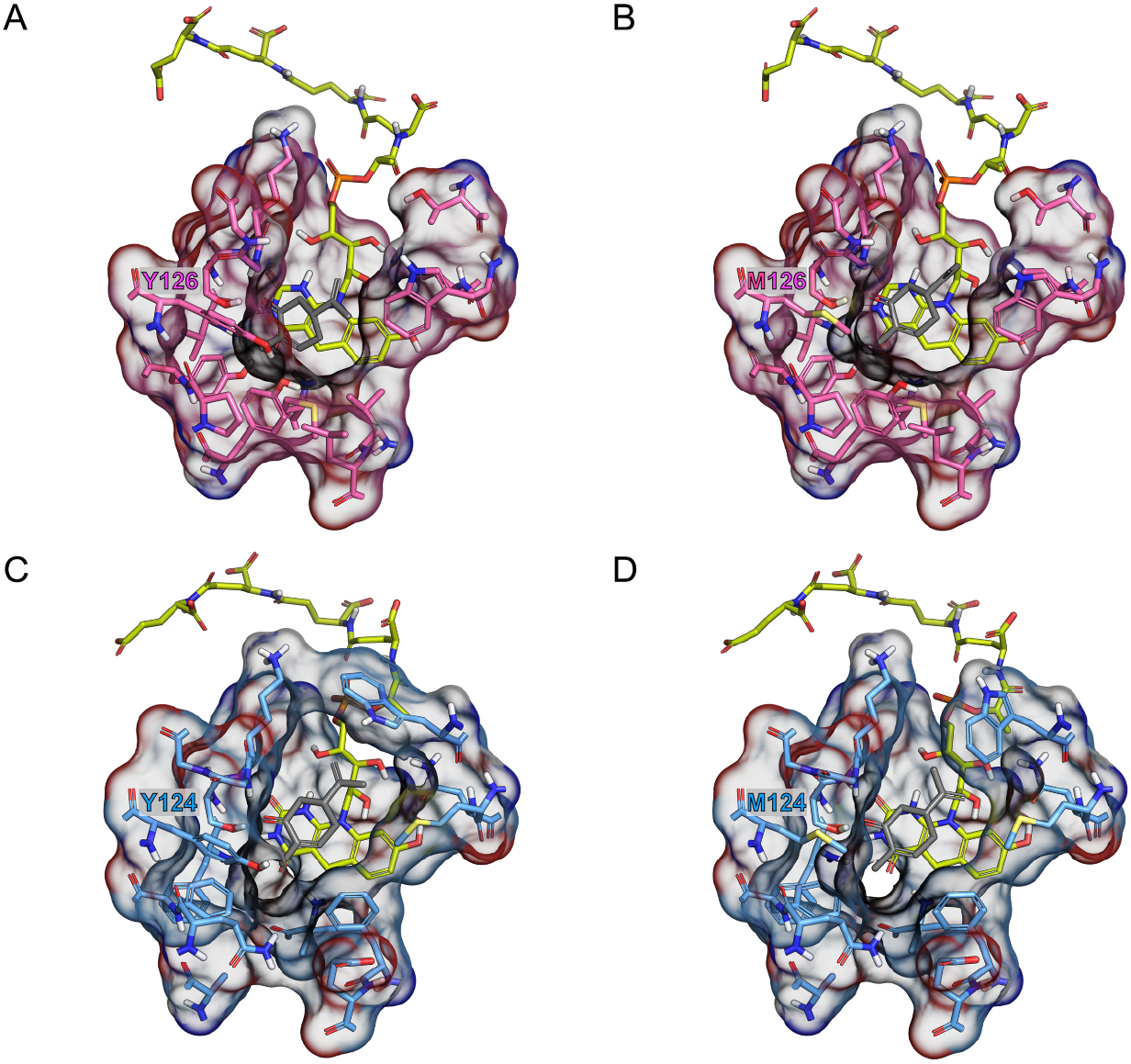
Induced fit docking poses of **1a** (gray) with wildtypes (A, C) and Y124/126M mutants (B, D). MSMEG_2850 and MSMEG_2027 variants are colored blue and pink, respectively. For both mutants the lowest energy pose has the same binding mode as the wildtype enzyme, suggesting that the altered stereochemical outcome with these mutants results from altered facial selectivity for the protonation step following hydride transfer.

We subsequently tested these variants with (*S*)-(+)-carvone (**2a**), the antipode of **1a**. The V30M/Y, V65L, and P122R variants of MSMEG_2027 had essentially identical selectivity for (2*S*)-**2b** (91.9–92.4% *de*, *cf*. 91.8% for WT), as well as comparable conversion (88.8–92.9%, *cf*. 88.4% wildtype) as shown in Figure 2B and Table S3. The lack of pronounced improvement compared to **1a** may reflect the greater selectivity of the wildtype with **2a** compared to **1a**. As before, MSMEG_2027-Y126M showed markedly altered selectivity, although it retained a preference for (2*S*)-**2b** with 28% *de*. Interestingly, conversion of **2a** was not attenuated in this mutant as it was with **1a**. It appears that V30, V65, or P122 are critical sites for altering the stereoselectivity of both carvone enantiomers. Y120Q/A and Y126M, on the other hand, displayed decreased stereoselectivity for the (2*S*)-configured product of both **2a** and **1a**, with only Y126M showing a weakly inverted preference for (2*R*)-**1b**.

MSMEG_2850-V53I and Q120Y showed increased selectivity for (2*S*)-**2b** (87.0 and 86.0%, *cf*. 83.0% *de* for wildtype), along with a moderate increase in conversion (76.1 and 87.2%, *cf*. 64.0% for wildtype) as shown in Figure 2B and Table S4. Selectivity was vastly reduced with Y124M, while F121M showed weakly switched stereoselectivity (−14.2% *de*). The F121M variant had the greatest impact on reversing stereoselectivity for **2a**, while the Y124M had the greatest impact on switching stereoselectivity for **1a**.

### Screening of variants against a panel of prochiral substrates

We expanded our investigation to include an array of prochiral substrates that we had previously tested against the wildtype enzymes (Figure 4). The values of conversions and stereoselectivity of **1a**–**10a** are shown in Tables S3 and S4. Individual plots showing performance of variants for each substrate are shown in Figures S4–S7.

**Figure 4.**
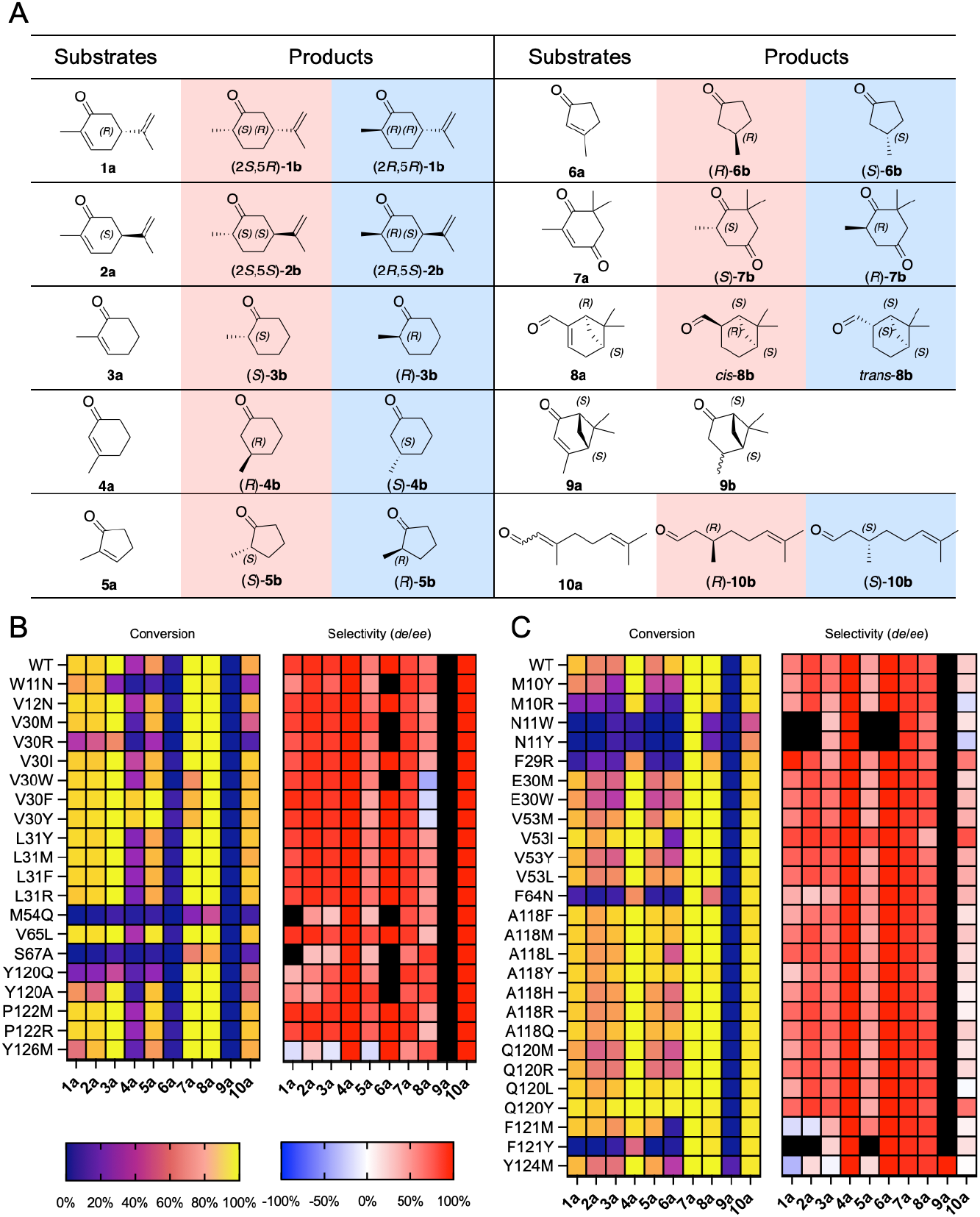
Activity and stereoselectivity of MSMEG_2027 and MSMEG_2850 variants against a panel of prochiral substrates. (A) Structures of substrates and products. Major products of wildtype enzymes are shaded red, minor products are shaded blue. Heatmaps were created to summarize the conversion and selectivity of MSMEG_2027 (B) and MSMEG_2850 (C) variants against substrates **1a**–**10a**. Positive values indicate selectivity that is similar to wildtypes, whereas negative values indicate reversed selectivity. Combinations where low conversion prevented accurate determination of selectivity are indicated in black.

### Cyclic enones bearing methyl substituents (3a-7a)

Compound **3a** is a substructure of both **1a** and **2a** and most variants showed a strong preference for the (*S*)-configured products, consistent with the expected similarity in binding orientation. MSMEG_2027-Y126M and MSMEG_2027-Y124M were again exceptions, showing weakly reversed selectivity (less than 20% *ee*, Figure 4, Tables S3 and S4). None of the MSMEG_2027 variants showed substantial improvements in conversion or selectivity compared to the wildtype. For MSMEG_2850, 15 mutants (V53I/Y/L/M, Q120Y/L/R, A118Q/L/F/R/H, E30M/W, and M10Y) showed improved selectivity for (*S*)-**3b**, with up to 85% *ee* obtained with V53I, compared to 62.9% with the wildtype. V53I also showed an increase in conversion (93% *cf*. 62.6% for the wildtype). Q120Y had the highest conversion (97.2%) and the second highest enantioselectivity (77.2% *ee*). The same effect of these two mutations was also observed with the carvones. Likewise, the MSMEG_2027 variants M54Q and S67A, as well as the N11W/Y, F64N and F121Y variants of MSMEG_2850 showed severely reduced activity compared to wildtype.

Contrastingly, reduction of **4a** produced exclusively the (*R*)-configured product for all the variants. Conversions with MSMEG_2027 were generally low, however mutation of V30 to Tyr or Phe resulted in substantially higher activity (89.5 and 88.3%, respectively), compared to the wildtype (30.9%). Most variants of MSMEG_2850 showed good conversion of **4a** as did the wildtype enzyme. The exceptions were again the N11W/Y, F64N and F121Y variants that were inactive against carvone.

The cyclopentenone derivatives (**5a** and **6a**) showed similar enantioselectivity but lower conversion compared to their cyclohexanone analogues. With **5a**, seven variants (Y120A, V30M, V65L, P122M/R, L31R, and V12N) of MSMEG_2027 increased enantioselectivity for (*S*)-**5b** (up to 86.7% *ee* with Y120A *cf*. 60.9% *ee* with the wildtype). Consistent with the other 2-methyl substituted compounds **1a**, **2a**, and **3a**, Y126M reversed the selectivity of **5a** to yield (*R*)-**5b** albeit with only 18.5% *ee*. All hydrophobic V30 variants (V30Y/F/W/M/I) increased conversion (>99% with V30Y/F) compared to 70.8% with the wildtype. In contrast, V30R significantly reduced conversion to 30%, while the corresponding variant MSMEG_2850-F29R also showed decreased conversion (56.7%). Ten variants (V53I, F121M, A118R/Y/F/L/H/Q, Q120L, and V53L) of MSMEG_2850 showed both improved enantioselectivity and conversion. Among them, V53I significantly improved both stereoselectivity and conversion to 79.4 and 93.7%, respectively, compared to 44.5 and 62.6% with the wildtype. Y124M decreased enantioselectivity to 17.8% *ee* but did not invert selectivity, unlike with **3a**.

As with **4a**, **6a** was exclusively converted to the (*R*)-enantiomer. MSMEG_2027 variants showed poor conversion of these substrates, between 0 and 7.0%. V30Y and V30F enhanced conversion to 7.0 and 5.7%, respectively, up from 3.6% with the wildtype. The Q120Y/L and A118Y variants of MSMEG_2850 improved conversion of **6a**, (up to 97.5% with Q120Y) compared to 85.9% with the wildtype. Consistent with the other 3-methyl substituted compound **4a**, the Y126M and Y124M mutations did not affect the stereoselectivity of the reduction of **6a**.

With ketoisophorone **7a**, 14 variants of MSMEG_2027 had improved selectivity for the (*S*)-configured product, although improvements were modest (89.7% *ee*, with P122R *cf*. 82.7% with the wildtype). Most of these mutants retained the high conversions seen with the wildtype. The Y126M mutant greatly diminished enantioselectivity (54.4% *ee*) as observed with the other 2-methyl substituted cyclic enones **1a**, **2a**, **3a** and **5a**. All variants of MSMEG_2850 showed >99% conversion and showed only modest variation in selectivity (85.1–91.7% *ee*, *cf*. 87% for the wildtype). Interestingly, the N11W and N11Y variants showed complete conversion of **7a** despite showing little or no activity with the previous substrates. In contrast to MSMEG_2027-Y126M, the MSMEG_2850-Y124M variant showed increased selectivity for (*S*)-**7b** rather than switching to (*R*)-**7b** (91.7% *ee*, *cf*. to 87.0% with the wildtype).

### a-pinene derivatives myrtenal (8a) and verbenone (9a)

With (*R*)-myrtenal **8a**, all variants of MSMEG_2027 showed >99% conversion, except W11N, S67A and M54Q. In contrast to the activity with the carvone enantiomers, the S67A mutation resulted in only a moderate decrease of conversion (75.4%), indicating that the hydrogen bonding with the sidechain of S67 is not critical for activity with **8a**. The S67A, Y120A/Q, Y126M, and W11N mutations increased diasteroeselectivity *cis*) between 72.0 and 85.7% *de*, up from 63.6% with the wildtype. On the other hand, V30Y/F/W reversed stereoselectivity towards favoring the *trans*isomer, albeit with only moderate selectivity (−16.9 to −41.1% *de*). The other V30 variants, V30M/R/I, slightly lowered the *de (cis*). For MSMEG_2850, most variants retained activity of above 90% conversion. 13 Mutants improved *de (cis*) up to 86.9 and 85% with Y124M and F121M, compared to 72.8% with the wildtype. Only the V53I variant showed substantially reduced selectivity (37%) and no variant produced predominantly *trans*-**8b**.

Like **8a**, (1 *S*,5*S*)-(-)-verbenone **9a** contains the α-pinene scaffold, however unlike **8a**, activity with **9a** could not be detected by GC/MS using wildtype MSMEG_2027 and MSMEG_2850. Given the improved activity against the previous substrates we tested to see if any variants showed improved activity against **9a**. Only one variant, MSMEG_2850-Y124M, showed modest conversion (10.0%). No other variants had detectable activity by GC/MS. Only one product peak was observed in the GC/MS trace, however we were unable to assign stereochemistry of the product without a suitable reference material. Few studies have reported on **9a** as a substrate for ene-reductases, however it has been shown that **9a** is not a substrate for the well characterized OYE homologues PETNR, YqjM, OYE2 or TOYE.^35^ Modest activity has also been reported with isopiperitenone reductase, a member of the FMN-independent plant short-chain dehydrogenase/reductase family, as well as an enzyme extracted from cultured cells of *Nicotiana tabacum.^8^*, ^36^ Based on the results with **4a**, (which is a substructure of **9a**) the most likely product would be (1 *S*,2*S*,5*S*)-*trans*-verbanone **9b** (Cβ in *trans*-**9b** has the same relative configuration as Cβ in (*R*)-**4b**, however the CIP priority is altered due to the bicyclic structure of **9b**). This is supported by poses obtained with induced fit docking (Figure S8).

### Citral (10a) and its constituent isomers

With citral **10a,** 13 variants of MSMEG_2027 (L31F/Y/R/M, V30I/F/Y/W, V65L, P122R/M, V12N, and Y126M) showed increased conversion. The highest conversion was seen with L31F with 87.4%, up from 76.2% with the wildtype. Conversion was lower with the W11N, M54Q and S67A variants, consistent with the previous substrates. In contrast, all MSMEG_2850 variants showed comparable conversion to the wildtype (>90%) except for N11W, N11Y and F29R (46.6, 66.5 and 86.3%, respectively). All variants of MSMEG_2027 had the same high selectivity of the wildtype (>99% *R*). In contrast, MSMEG_2850 variants showed a wide range of selectivity. Nine variants (V53I/L/Y/M, Q120Y, F29R, F64N, E30W, and A118M) increased *ee* (*R*) compared to the wildtype (up to 77.5% with V53I, compared to 25% with the wildtype). Q120L, M10R, and N11Y switched selectivity to (*S*)-**10b** with −3.0, −17.1 and −24.1% *ee*, respectively.

**10a** occurs naturally as a 6:4 mixture of *cis*- and *trans*-isomers. Using pure samples of each isomer, we found previously that wildtype MSMEG_2027 reduced both isomers to (*R*)-**10b**, consistent with the high selectivity observed with the mixture.^10^ In contrast, wildtype MSMEG_2850 reduces *cis*-**10a** exclusively to (*R*)-**10b**, while *trans-* **10a** is reduced to (*S*)-**10b** with only 52.3% *ee*.^10^ We assayed selected variants with both isomers to gain further insight into the effects of these mutations (Figure S7B and C). The three variants (V53I, Q120Y, and F29R) with the highest selectivity for (*R*)- **10b** converted *cis-***10a** exclusively to (*R*)-**10b** as did the wildtype. These variants switched selectivity of *trans-**10a*** reduction to from (*S*)- to (*R*)-**10b** with 67.1, 49.9 and 26.6% *ee*, respectively (Table S4).

Similarly, the three variants (Q120L, M10R, and N11Y) that most strongly favored (*S*)-**10b**, did so by increasing selectivity for (*S*)-**10b** with *trans-**10a*** (−74.9, −70.7 and −66.8% respectively). With *cis*-**10a**, Q120L and M10R retained selectivity for (*R*)-**10b** with conversions of >99 and 97.2%, respectively. However, N11Y showed significantly reduced conversion of *cis*-**10a** (44.8%) along with slightly decreased *ee* (83.8%). Therefore, the higher selectivity for (*S*)-**10b** from **10a** with N11Y compared to M10R and Q120L results largely from reduced turnover of *trans*-**10a**, which yields predominantly (*R*)-**10b**, rather than greatly improved selectivity for (*S*)-**10b** with *trans-* **10a**.

### Effect of mutations on stereoselectivity

To gain a broader overview of the effects of each mutation on different substrates we calculated Spearman rank correlation coefficients between the selectivity (*de* or *ee*) between each pair of substrates. The results in Figure 5, Tables S5 and S6 show that for both enzymes the selectivity with 2-methyl substituted cyclic enones (**1a**, **2a**, **3a** and **5a**) are positively correlated with each other, although these correlations were not always statistically significant (Tables S5 and S6). This indicates that mutations that improve or invert selectivity with one substrate in this set are likely to have a similar effect on another substrate. In contrast, selectivity of **8a** was negatively correlated with that of **1a**, **2a**, **3a**, and **5a**, indicating that mutations that increased selectivity for *cis-* **8b** were associated with lower (*S*)-selectivity with these substrates. However, mutations that strongly affect selectivity with **8a** show selectivity similar to that of the wildtype with these substrates and vice-versa, indicating that the selectivity between these sets of substrates may be orthogonal (Figures S9 and S10). Substrates **4a**, **6a**, **9a** showed no variation in selectivity, nor did **10a** with MSMEG_2027, and therefore Spearman correlations could not be calculated in these cases.

**Figure 5.**
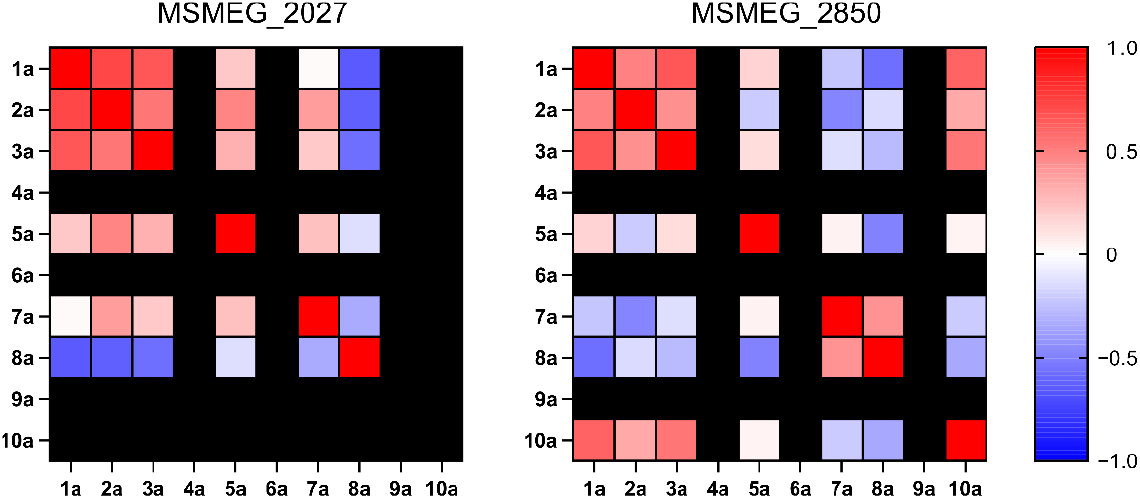
Spearman correlation coefficients between the effect of mutations on selectivity of different substrates. For substrates that did not show any difference between variants no correlation could be calculated, and these are shown as black squares. Tabulated data with *p* values for MSMEG_2027 and MSMEG_2850 are listed in Table S5 and S6, respectively.

Interestingly, MSMEG_2027 and MSMEG_2850 variants showed markedly different correlations between selectivity with **7a** and that of the other 2-methyl cyclic enones. With MSMEG_2027 **7a** has weak positive correlations with these substrates, while with MSMEG_2850 these are weakly negative, although in most cases these are not statistically significant (Table S6).

The mutations that most affected stereoselectivity without substantial loss of activity were examined for each substrate (Figure 6). In most cases, the mutants with improved wildtype selectivity also showed increased conversion of substrate. For MSMEG_2027, a mutation at V30 in loop 1 affected the stereoselectivity most frequently. This position might therefore be considered as a hot spot. Against most substrates, the V30Y mutation had the most improved conversion rates (Figure 6A). Met replacement at Y126 in α4 reversed the selectivity for 2-methyl substituted enones. For MSMEG_2850, an Ile replacement at V53 in loop 3 had the greatest effects on the stereoselectivity for most of the tested substrates (Figure 6B). V53 could thus also be considered a hot spot for MSMEG_2850. Mutations at Q120 in loop 10 also showed a dramatic impact with some substrates, most notably Q120L reversing the selectivity with trans-**10a**. The most significant improvement in conversion rate was achieved by mutation of Q120 to Tyr. As with Y126M, Y124M in α4 reversed the stereoselectivity for 2-methyl substituted enones. Only Y124M displayed moderate conversion, whereas no other variants showed detectable activity.

**Figure 6.**
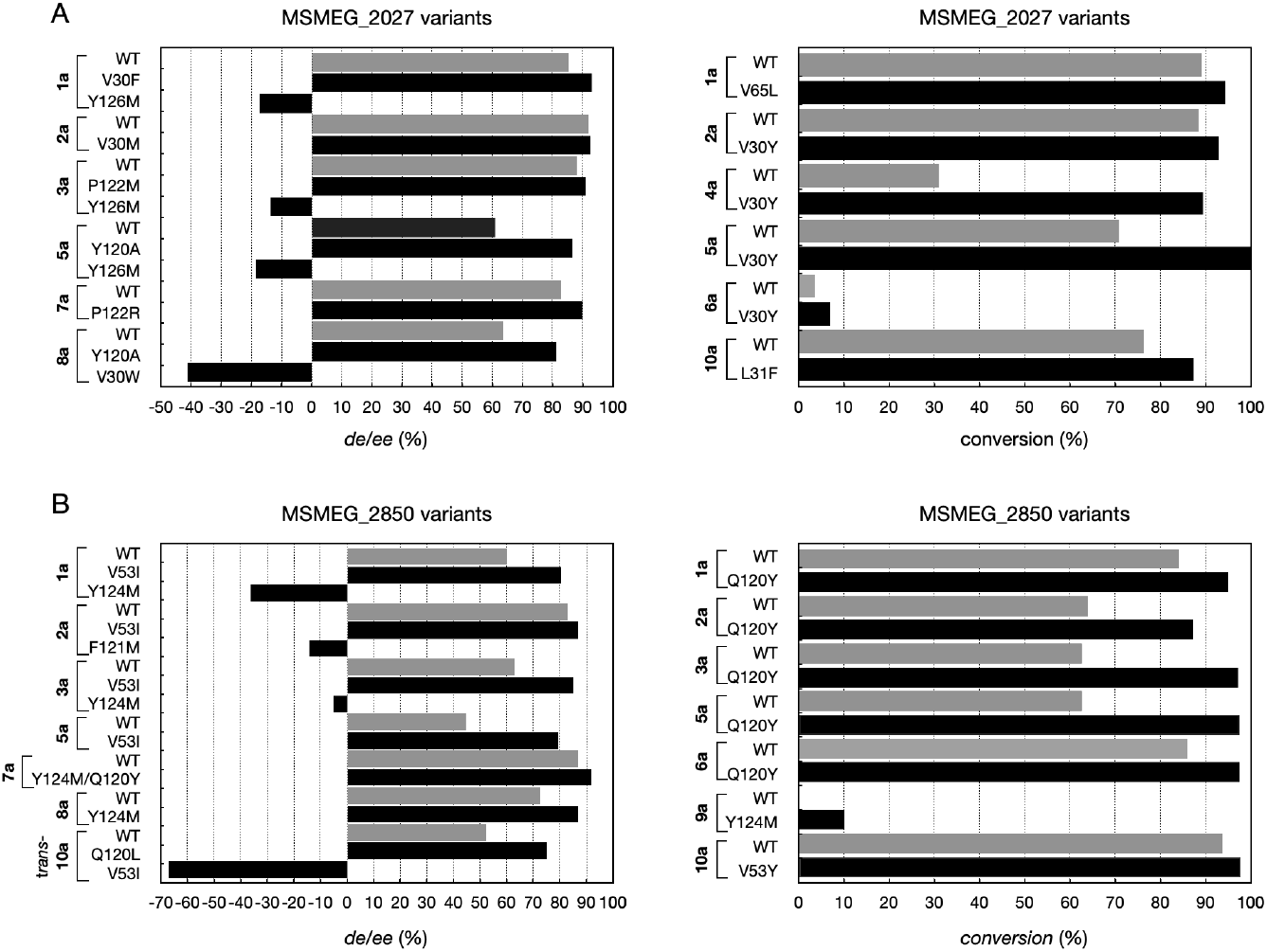
Mutants of MSMEG_2027 (A) and MSMEG_2850 (B) that most affected stereoselectivity and conversion. Mutants showing the greatest improvement in conversion and *de/ee* or the greatest reverse in *de/ee* are selected. The substrates against which wildtype (WT) has already achieved >99% *ee*/*de* or conversion were not shown. Wildtype *de*/*ee* values are shown in positive and reversed *de*/*ee* values are shown in negative.

### Computational docking and relative free energy analysis

To gain insight into the effects of these mutations we performed molecular docking analysis along with MM-GBSA to calculate the differences between wildtype and mutant proteins in binding free energy with the substrates. MSMEG_2027 mutations at V30 mostly impacted stereoselectivity and conversion, which suggests that V30 might be a “hot spot” that has a large effect on activity/specificity. Intriguingly, V30 is unique to MSMEG_2027 amongst the currently characterized FDOR-A enzymes that generally have Phe at this position (Figure 1C). In wildtype MSMEG_2027, the short side chain of V30 only forms weak hydrophobic interactions with the substrates (Figure 7A and C), while V30F and V30M can form stronger interactions with the substrates, reducing binding free energy (Figure 7B and D). Enhanced stereoselectivity and conversion can be explained by these lower binding energies with substrates in the same binding pose as the wildtype. We can expect a similar effect on other substrates with the hydrophobic mutations at V30 (V30Y/F/W/M/I) that improved conversion and selectivity (Figure 4 and S11D). It is interesting to note that V30W switched the stereoselectivity of **8a** despite the reduced binding energy with **8a** in the same orientation as the wildtype (Figure 7E and F). This will be discussed later in the section.

In wildtype MSMEG_2850 the methyl group at Cα of **1a** did not form any interaction with the short side chain of V53 (Figure 7G), however hydrophobic interactions are formed after mutation to Ile, leading to higher binding affinity (Figure 7H). Induced fit docking (IFD) confirms that the V53I variant can establish hydrophobic interactions with other 2-methyl substituted substrates (**2a**, **3a**, and **5a**) as well (Figure S12A-F). These are consistent with the increased stereoselectivity and conversion rates that were experimentally observed in the V53I variant.

**Figure 7.**
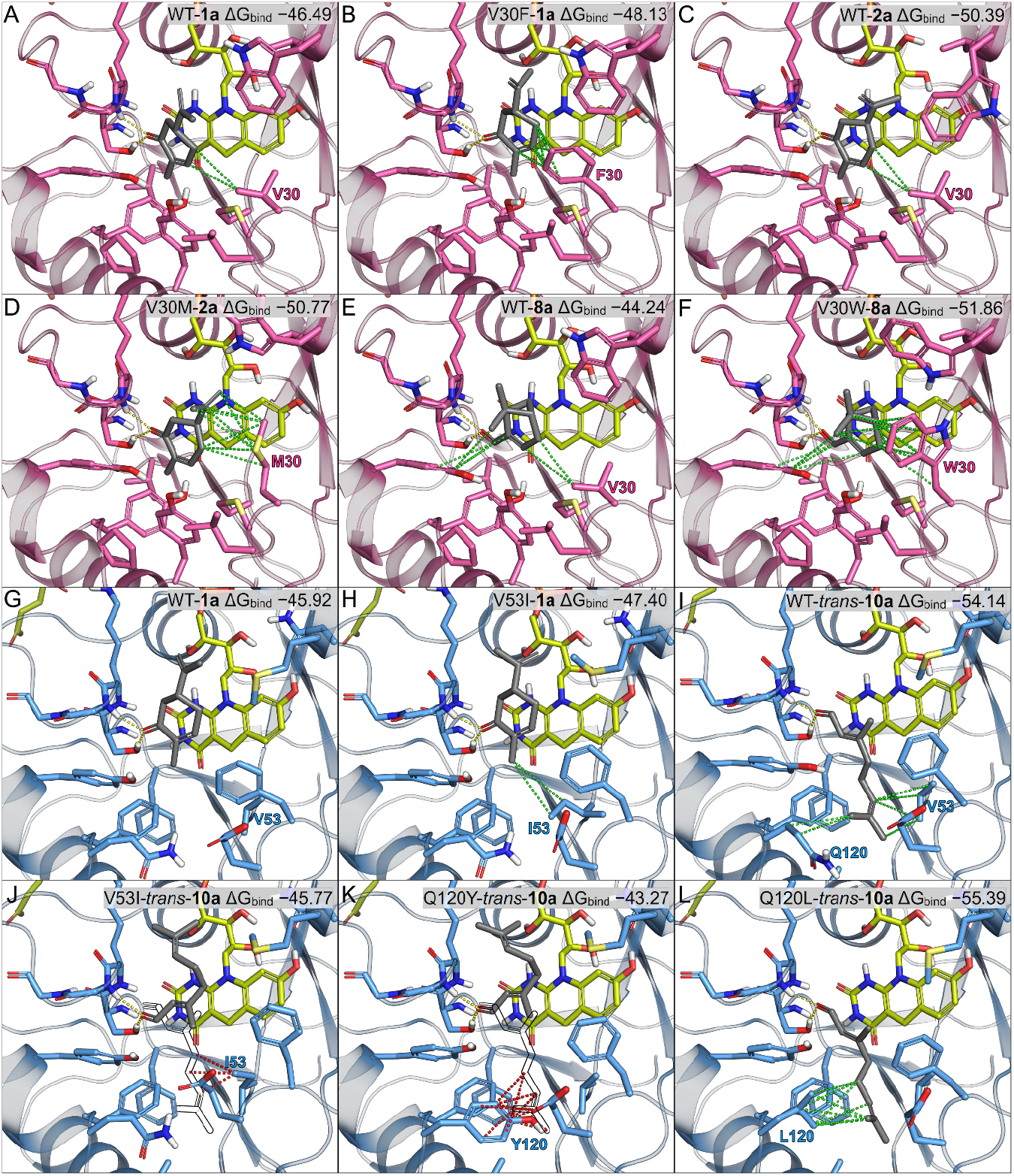
Induced fit docking (IFD) poses and MM-GBSA (Δ*G*bi⊓d, kcal mol^-1^) of wildtype FDORs and mutants at hot spot. Hydrogen bonds are represented by yellow dashed lines, and relevant hydrophobic interactions by green dashed lines. Steric clashes are indicated by red dashed lines with an imaginary wildtype binding pose of *trans*-**10a** (black outline) (J and K).

In the wildtype *trans-**10a*** can bind with the bulky Cβ substituent in the cavity created by V53 and Q120, forming a hydrophobic interaction (Figure 7I). However, in the V53I and Q120Y mutants this cavity is constricted, thereby preventing *trans-**10a*** binding in the same orientation due to a steric clash (Figure 7J and K). This necessitates a flipped binding mode that in turn leads to the observed reversed stereoselectivity. On the other hand, a Leu replacement at position 120 can form additional hydrophobic interactions with the tail of geranial in the classical binding mode, increasing stereoselectivity (Figure 7L).

We identified several mutations that resulted in a switch of the stereochemical outcome of substrate reduction. Notably MSMEG_2027-Y126M and MSMEG_2850- Y124M show reversed selectivity for cyclic enones bearing a substituent on Cα, but not cyclic enones substituted on Cβ. The stereochemistry of Cβ is solely determined by the binding mode adopted by the substrate, whereas Cα is susceptible to changes in both binding mode and hydrogenation mode.^25^ Induced fit docking results show that the most favored binding mode of these substrates is not “flipped” (Figure 8, S11, and S12), in contrast to many stereocomplementary OYE variants reported in the literature, and corroborates the hypothesis that the hydrogenation mode is perturbed towards favoring *syn* relative to *anti* addition.

**Figure 8.**
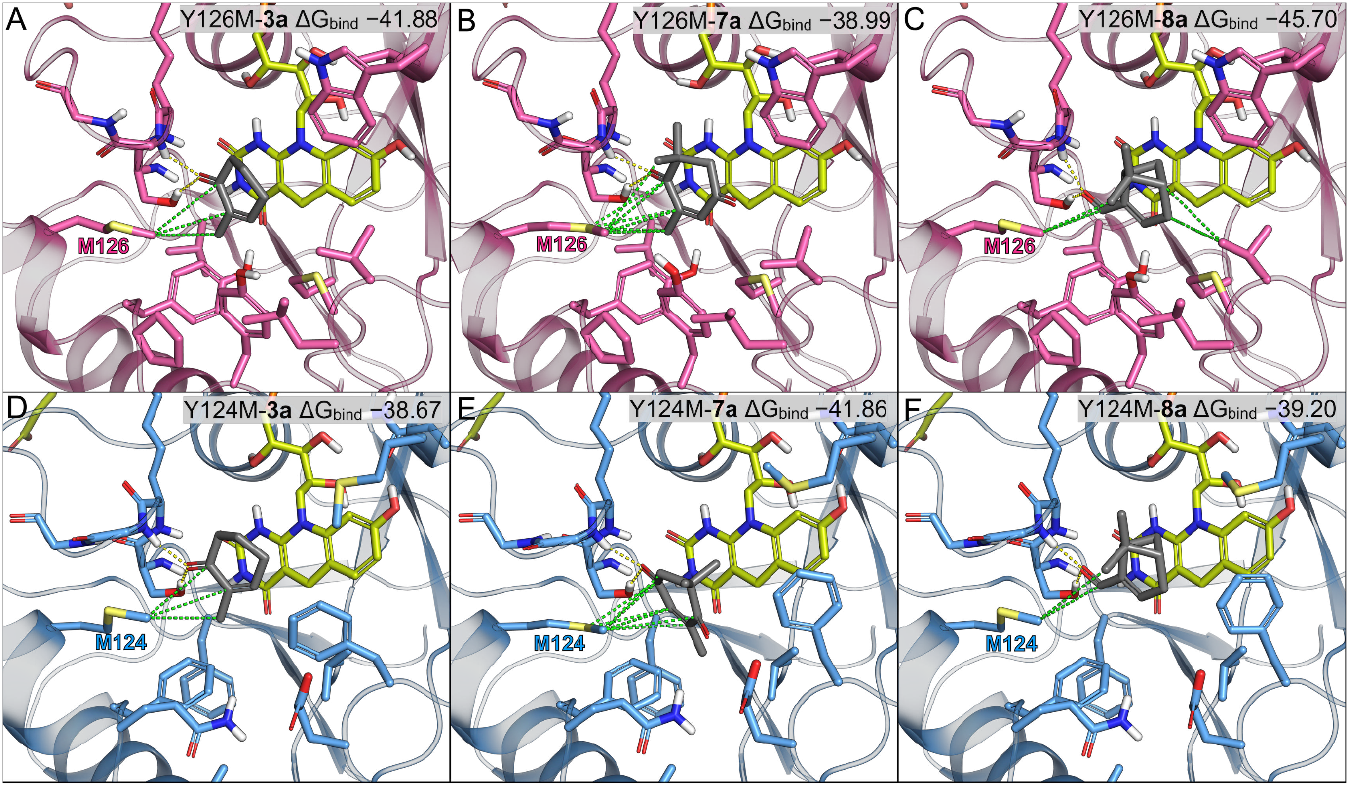
Induced fit docking (IFD) poses and MM-GBSA (Δ*G*_bind_, kcal mol^-1^) of mutants that switch stereoselectivity against 2-methyl substituted alkenes (**1a**, **2a**, **3a**, and **5a**). MSMEG_2850 and MSMEG_2027 variants are colored blue and pink, respectively. Hydrogen bonds are represented by yellow dashed lines, and relevant hydrophobic interactions by green dashed lines.

A notable difference between these two variants is their selectivity with **7a**. This compound has two carbonyl groups that could bind in the oxyanion hole, and we previously obtained docking poses of both orientations with the wildtype enzymes^.10^ With the C4-carbonyl as the activating group protonation occurs at unsubstituted C3, whereas with the C1-carbonyl as the activating group protonation take place at methyl-substituted C2. Through net *anti* addition of H_2_, both poses would yield the observed major product (*S*)-**7b**. Because of this there is ambiguity about which carbonyl serves as the activating group. However, each pose should be affected differently by the Y124M or Y126M mutations: with C4 as the activating group the stereochemical outcome is not affected by which face protonation occurs from, whereas with C1 as the activating group proton transfer determines the stereochemistry. Only Y126M affected stereoselectivity, indicating the C1-carbonyl binds in the oxyanion hole in MSMEG_2027-Y126M and the C4-carbonyl binds in the oxyanion hole in MSMEG_2850-Y124M. These conclusions were supported by IFD analysis (Figure S11H and S12I). Interestingly, these binding modes are consistent with the results with **4a** and **3a**, where MSMEG_2027 showed higher activity with the 2-methyl substituent, while MSMEG_2850 preferred the 3-methyl substitution. This is also consistent with the Spearman correlation between **7a** and other 2-methyl substituted substrates being positive for MSMEG_2027 but negative for MSMEG_2850 (Figure 5).

In MSMEG_2850, F29 and V53 seem to be key resides for the preference of 3- methyl substituted substrate. The small side chain of V53 allows space for the methyl group at C3 and F29 can form hydrophobic interactions with the methyl group, further lowering the binding free energy (Figure S13A and B). Mathew *et al*., previously reported that FDR-Mha, FDR-Rh1, and FDR-Rh2 also prefer 3-methyl cyclic enones over 2-methyl cyclic enones.^9^ These enzymes have Phe at the equivalent position of F29 and either Val or the slightly larger Leu at the equivalent of V53 (Figure 1C). In MSMEG_2027, M54 forms a hydrophobic interaction with the methyl group at C2 but causes steric hindrance with a methyl group at C3, which explains the preference for substrates bearing a methyl group at C2 over C3 (Figure S13C and D). Unlike F29 in MSMEG_2850, the short side chain of V30 cannot directly interact with the substrates. These explanations are supported by the behavior of the mutants V30F and V53M. The V30F variant of MSMEG_2027 increased the conversion rate of the 3-methyl substituted substrate **4a** and the V53M variant of MSMEG_2850 improved the stereoselectivity and conversion rate of the 2-methyl substituted substrate **3a** (Tables S3 and S4).

The tyrosine at position 126 (MSMEG_2027) is highly conserved in FDOR-As and, given its position in the active site, might be assumed to be mechanistically analogous to the highly conserved tyrosine residue in the active sites of OYE homologues (Y196 in OYE). In OYE homologues this residue is thought to be the proton donor given that the Y196F variant of OYE is incapable of turning over 2-cyclohexenone, and QM-MM calculations with YqjM support this role.^37, 38^ Loss of the tyrosine hydroxyl leading to inhibited *anti* addition of H_2_ would account for the observed change in selectivity in the Y126M variants of the FDORs. Importantly however, these variants retain considerable activity with many substrates, including ones that are poorly turned over by the wildtype enzymes. Furthermore, QM-MM calculations on the activation of pretomanid by Ddn indicate that proton transfer from tyrosine, either directly or mediated through a solvent molecule, is energetically unfavorable and that hydronium ions in the solvent are more likely to be the proton donor.^39^ Indeed, the equivalent mutant of Ddn (Y123M) retains ~75% of the wildtype activity with menadione and delamanid, although it completely lost activity with pretomanid.^18^ This same study found that aromatic residues at this position were important for creating a hydrophobic shield for the substrate against solvent during hydride transfer, as aliphatic residues at this position lead to loss of activity with pretomanid.^18, 40^ Disruption of this hydrophobic shield may therefore be the underlying cause of the proposed altered hydrogenation mode. This also explains the effect of the F121M mutation in MSMEG_2850.

Interestingly, selectivity with **8a** is largely unaffected by mutation at this conserved tyrosine. Both wildtype enzymes favor *cis*-**8b**, which can only form through *syn* addition of H_2_ if the docking poses reflect the Michaelis complex (Figure 7E and S12J). Mutations to V30 of MSMEG_2027 that increase the steric bulk shift selectivity towards *trans*-**8b** but this shift is particularly strong with aromatic residues as this position. Potentially these aromatic substitutions could extend the hydrophobic shield and thereby affect protonation mechanism, although the equivalent position on MSMEG_2850 is F29 and has higher selectivity for *cis*-**8b** than wildtype MSMEG_2027. In our substrate panel, **8a** is unique in that it contains an endocyclic C=C bond with an activating group external to the ring; these features may influence the hydrogenation mode. Additionally, the bicyclic ring system of **8a** is observed to be tilted away from the F_420_ cofactor when compared to other substrates with planar ring systems. This structural characteristic may facilitate the transfer of hydride and proton on the same side. Investigating the selectivity of other common OYE substrates that have these features such as perillaldehyde may be informative in this respect.

The mode of hydrogenation has been unambiguously determined in several ene-reductases through the use of isotope labelling experiments with examination of the products by ^1^H and/or ^2^H NMR.^25, 36^ The two hydrogens at C4 of NAD(P)H are enantiotopic and both may be selectively labelled by choosing an appropriate reductase.^41^ Similarly, the hydrogens at C5 of F_420_H_2_ are enantiotopic and can be labelled analogously.^42^ The FDORs transfer the hydride from the *re*-face of F_420_H_2_, however all F_420_ reductases described to date are *si*-face selective, thereby prohibiting this approach with the FDORs.^43^ An alternative method is to follow the stereochemical outcome of cyclic tetrasubstituted alkenes. Using α-methylketoisophorone Venturi *et al*., have shown that FDR-Rh1, FDR-Rh2 and FDR-Mha all favor *anti* addition, at least with this substrate.^44^

### Measurement of Michaelis–Menten enzyme kinetics

For a more detailed analysis, we measured apparent steady state kinetic parameters for selected variants with the nine substrates (Tables 1 and 2). Michaelis–Menten plots are shown in Figures S14 and S15. Kinetic parameters provided more detailed information about catalytic activity, particularly in case where conversions between variants are not distinctive; for example, most variants showing >99% conversion from GC/MS analysis.

**Table 1.** Kinetics parameters for MSMEG_2027 variants.

|  | Parameter | Wildtype | V30Y | V65L | P122M |
| --- | --- | --- | --- | --- | --- |
| **1a** | $K_M$ (mM) | $1.29 \pm 0.32$ | $0.61 \pm 0.06$ | $0.79 \pm 0.12$ | $0.85 \pm 0.19$ |
| | $k_{cat}$ ($s^{-1}$) | $0.11 \pm 0.008$ | $0.05 \pm 0.001$ | $0.12 \pm 0.005$ | $0.11 \pm 0.007$ |
| | $k_{cat}/K_M$ ($M^{-1} s^{-1}$) | $(8.21 \pm 1.53) \times 10^1$ | $(7.54 \pm 0.63) \times 10^1$ | $(1.53 \pm 0.19) \times 10^2$ | $(1.26 \pm 0.23) \times 10^2$ |
| | $K_i$ (mM) | — | — | — | — |
| **2a** | $K_M$ (mM) | $1.30 \pm 0.28$ | $0.51 \pm 0.06$ | $1.00 \pm 0.15$ | $0.88 \pm 0.19$ |
| | $k_{cat}$ ($s^{-1}$) | $0.09 \pm 0.006$ | $0.06 \pm 0.002$ | $0.10 \pm 0.005$ | $0.09 \pm 0.005$ |
| | $k_{cat}/K_M$ ($M^{-1} s^{-1}$) | $(6.86 \pm 1.121) \times 10^1$ | $(1.17 \pm 0.109) \times 10^2$ | $(1.02 \pm 0.117) \times 10^2$ | $(9.72 \pm 1.62) \times 10^1$ |
| | $K_i$ (mM) | — | — | — | — |
| **3a** | $K_M$ (mM) | $2.56 \pm 0.48$ | $1.63 \pm 0.10$ | $1.64 \pm 0.37$ | $3.96 \pm 0.88$ |
| | $k_{cat}$ ($s^{-1}$) | $0.62 \pm 0.05$ | $0.74 \pm 0.016$ | $0.39 \pm 0.03$ | $0.84 \pm 0.08$ |
| | $k_{cat}/K_M$ ($M^{-1} s^{-1}$) | $(2.43 \pm 0.31) \times 10^2$ | $(4.54 \pm 0.20) \times 10^2$ | $(2.38 \pm 0.40) \times 10^2$ | $(2.13 \pm 0.29) \times 10^2$ |
| | $K_i$ (mM) | — | — | — | — |
| **4a** | $K_M$ (mM) | $4.29 \pm 3.46$ | $3.33 \pm 0.53$ | $19.5 \pm 19.6$ | $4.46 \pm 3.29$ |
| | $k_{cat}$ ($s^{-1}$) | $1.10 \pm 0.41$ | $1.04 \pm 0.11$ | $2.25 \pm 1.63$ | $1.13 \pm 0.39$ |
| | $k_{cat}/K_M$ ($M^{-1} s^{-1}$) | $(2.57 \pm 1.23) \times 10^2$ | $(3.11 \pm 0.17) \times 10^2$ | $(1.16 \pm 0.34) \times 10^2$ | $(2.53 \pm 1.09) \times 10^2$ |
| | $K_i$ (mM) | — | $14.3 \pm 3.8$ | — | — |
| **5a** | $K_M$ (mM) | $1.00 \pm 0.13$ | $0.67 \pm 0.13$ | $1.32 \pm 0.22$ | $1.32 \pm 0.18$ |
| | $k_{cat}$ ($s^{-1}$) | $0.55 \pm 0.02$ | $0.63 \pm 0.04$ | $0.62 \pm 0.03$ | $0.67 \pm 0.03$ |
| | $k_{cat}/K_M$ ($M^{-1} s^{-1}$) | $(5.50 \pm 0.54) \times 10^2$ | $(9.35 \pm 1.50) \times 10^2$ | $(4.69 \pm 0.60) \times 10^2$ | $(5.08 \pm 0.53) \times 10^2$ |
| | $K_i$ (mM) | — | — | — | — |
| **6a** | $K_M$ (mM) | $0.12 \pm 0.05$ | $0.03 \pm 0.011$ | — | $0.03 \pm 0.007$ |
| | $k_{cat}$ ($s^{-1}$) | $0.01 \pm 0.001$ | $0.01 \pm 0.001$ | — | $0.01 \pm 0.001$ |
| | $k_{cat}/K_M$ ($M^{-1} s^{-1}$) | $(1.16 \pm 0.41) \times 10^2$ | $(2.99 \pm 1.11) \times 10^2$ | — | $(2.34 \pm 0.60) \times 10^2$ |
| | $K_i$ (mM) | — | — | — | — |
| **7a** | $K_M$ (mM) | $3.88 \pm 0.68$ | $0.18 \pm 0.02$ | $2.51 \pm 0.22$ | $2.20 \pm 0.15$ |
| | $k_{cat}$ ($s^{-1}$) | $0.39 \pm 0.05$ | $0.01 \pm 0.001$ | $0.45 \pm 0.02$ | $0.26 \pm 0.007$ |
| | $k_{cat}/K_M$ ($M^{-1} s^{-1}$) | $(1.00 \pm 0.06) \times 10^2$ | $(4.28 \pm 0.41) \times 10^1$ | $(1.78 \pm 0.06) \times 10^2$ | $(1.18 \pm 0.06) \times 10^2$ |
| | $K_i$ (mM) | $26.9 \pm 11.4$ | — | $69.9 \pm 28.3$ | — |
| **8a** | $K_M$ (mM) | $0.31 \pm 0.03$ | $0.30 \pm 0.06$ | $0.50 \pm 0.03$ | $0.49 \pm 0.02$ |
| | $k_{cat}$ ($s^{-1}$) | $0.74 \pm 0.017$ | $1.03 \pm 0.07$ | $0.83 \pm 0.014$ | $0.82 \pm 0.011$ |
| | $k_{cat}/K_M$ ($M^{-1} s^{-1}$) | $(2.39 \pm 0.196) \times 10^3$ | $(3.48 \pm 0.46) \times 10^3$ | $(1.64 \pm 0.09) \times 10^3$ | $(1.69 \pm 0.07) \times 10^3$ |
| | $K_i$ (mM) | — | $52.6 \pm 31.4$ | — | — |
| **10a** | $K_M$ (mM) | $0.67 \pm 0.09$ | $0.43 \pm 0.09$ | $0.59 \pm 0.06$ | $0.54 \pm 0.07$ |
| | $k_{cat}$ ($s^{-1}$) | $0.14 \pm 0.005$ | $0.10 \pm 0.009$ | $0.14 \pm 0.004$ | $0.13 \pm 0.005$ |
| | $k_{cat}/K_M$ ($M^{-1} s^{-1}$) | $(2.12 \pm 0.24) \times 10^2$ | $(2.46 \pm 0.35) \times 10^2$ | $(2.38 \pm 0.19) \times 10^2$ | $(2.44 \pm 0.27) \times 10^2$ |
| | $K_i$ (mM) | — | $23.4 \pm 9.8$ | — | — |

**Table 2.** Kinetics parameters for MSMEG_2027 variants.

|  | Parameter | Wildtype | V53I | Q120Y | F121M | Y124M |
| --- | --- | --- | --- | --- | --- | --- |
| **1a** | $K_M$ (mM) | $1.62 \pm 0.18$ | $0.99 \pm 0.11$ | $0.51 \pm 0.10$ | $1.62 \pm 0.11$ | $2.18 \pm 0.43$ |
| | $k_{cat}$ ($s^{-1}$) | $0.08 \pm 0.003$ | $0.08 \pm 0.003$ | $0.06 \pm 0.003$ | $0.08 \pm 0.002$ | $0.07 \pm 0.005$ |
| | $k_{cat}/K_M$ ($M^{-1} s^{-1}$) | $(5.10 \pm 0.41) \times 10^1$ | $(8.09 \pm 0.72) \times 10^1$ | $(1.28 \pm 0.22) \times 10^2$ | $(4.63 \pm 0.22) \times 10^1$ | $(3.18 \pm 0.44) \times 10^1$ |
| | $K_i$ (mM) | — | — | — | — | — |
| **2a** | $K_M$ (mM) | $2.07 \pm 0.39$ | $1.62 \pm 0.33$ | $0.58 \pm 0.12$ | $0.71 \pm 0.13$ | $1.61 \pm 0.33$ |
| | $k_{cat}$ ($s^{-1}$) | $0.05 \pm 0.004$ | $0.03 \pm 0.002$ | $0.03 \pm 0.002$ | $0.03 \pm 0.002$ | $0.03 \pm 0.002$ |
| | $k_{cat}/K_M$ ($M^{-1} s^{-1}$) | $(2.42 \pm 0.32) \times 10^1$ | $(1.65 \pm 0.24) \times 10^1$ | $(5.20 \pm 0.86) \times 10^1$ | $(4.07 \pm 0.60) \times 10^1$ | $(1.80 \pm 0.27) \times 10^1$ |
| | $K_i$ (mM) | — | — | — | — | — |
| **3a** | $K_M$ (mM) | $9.18 \pm 2.22$ | $6.60 \pm 3.78$ | $4.00 \pm 2.45$ | $3.45 \pm 0.83$ | $5.19 \pm 2.68$ |
| | $k_{cat}$ ($s^{-1}$) | $1.59 \pm 0.23$ | $1.63 \pm 0.49$ | $1.25 \pm 0.34$ | $1.04 \pm 0.11$ | $1.38 \pm 0.35$ |
| | $k_{cat}/K_M$ ($M^{-1} s^{-1}$) | $(1.73 \pm 0.19) \times 10^2$ | $(2.48 \pm 0.73) \times 10^2$ | $(3.11 \pm 1.15) \times 10^2$ | $(3.02 \pm 0.45) \times 10^2$ | $(2.66 \pm 0.76) \times 10^2$ |
| | $K_i$ (mM) | — | — | — | — | — |
| **4a** | $K_M$ (mM) | $6.73 \pm 6.84$ | $2.37 \pm 1.37$ | $4.54 \pm 5.75$ | $2.14 \pm 0.45$ | $4.04 \pm 2.10$ |
| | $k_{cat}$ ($s^{-1}$) | $4.02 \pm 3.30$ | $1.47 \pm 0.56$ | $2.29 \pm 2.31$ | $1.15 \pm 0.15$ | $2.25 \pm 0.84$ |
| | $k_{cat}/K_M$ ($M^{-1} s^{-1}$) | $(5.97 \pm 1.28) \times 10^2$ | $(6.20 \pm 1.37) \times 10^2$ | $(5.05 \pm 1.45) \times 10^2$ | $(5.38 \pm 0.49) \times 10^2$ | $(5.58 \pm 0.89) \times 10^2$ |
| | $K_i$ (mM) | $4.4 \pm 5.3$ | $9.2 \pm 7.3$ | $2.7 \pm 3.8$ | $17.6 \pm 6.7$ | $9.2 \pm 6.7$ |
| **5a** | $K_M$ (mM) | $7.67 \pm 3.21$ | $5.800 \pm 1.67$ | $6.27 \pm 1.33$ | $1.75 \pm 0.78$ | $0.59 \pm 0.18$ |
| | $k_{\text{cat}}$ ($\text{s}^{-1}$) | $0.16 \pm 0.04$ | $0.19 \pm 0.03$ | $0.18 \pm 0.019$ | $0.017 \pm 0.003$ | $0.07 \pm 0.005$ |
| | $k_{\text{cat}}/K_{\text{M}}$ ($\text{M}^{-1} \text{s}^{-1}$) | $(2.09 \pm 0.42) \times 10^1$ | $(3.31 \pm 0.51) \times 10^1$ | $(2.81 \pm 0.31) \times 10^1$ | $(9.49 \pm 3.05) \times 10^0$ | $(1.11 \pm 0.28) \times 10^2$ |
| | $K_{\text{i}}$ (mM) | — | — | — | — | — |
| **6a** | $K_{\text{M}}$ (mM) | $4.30 \pm 0.74$ | $0.60 \pm 0.25$ | $1.03 \pm 0.26$ | $0.02 \pm 0.006$ | $0.09 \pm 0.03$ |
| | $k_{\text{cat}}$ ($\text{s}^{-1}$) | $0.10 \pm 0.008$ | $0.01 \pm 0.001$ | $0.02 \pm 0.001$ | $0.003 \pm 0.0001$ | $0.01 \pm 0.0004$ |
| | $k_{\text{cat}}/K_{\text{M}}$ ($\text{M}^{-1} \text{s}^{-1}$) | $(2.29 \pm 0.23) \times 10^1$ | $(8.73 \pm 2.90) \times 10^0$ | $(1.47 \pm 0.29) \times 10^1$ | $(1.65 \pm 0.49) \times 10^2$ | $(9.58 \pm 2.73) \times 10^1$ |
| | $K_{\text{i}}$ (mM) | — | — | — | — | — |
| **7a** | $K_{\text{M}}$ (mM) | $0.10 \pm 0.008$ | $0.29 \pm 0.030$ | $0.21 \pm 0.020$ | $3.93 \pm 0.18$ | $0.058 \pm 0.006$ |
| | $k_{\text{cat}}$ ($\text{s}^{-1}$) | $1.35 \pm 0.028$ | $0.51 \pm 0.013$ | $0.82 \pm 0.025$ | $1.05 \pm 0.02$ | $1.14 \pm 0.023$ |
| | $k_{\text{cat}}/K_{\text{M}}$ ($\text{M}^{-1} \text{s}^{-1}$) | $(1.35 \pm 0.09) \times 10^4$ | $(1.79 \pm 0.16) \times 10^3$ | $(3.92 \pm 0.29) \times 10^3$ | $(2.67 \pm 0.07) \times 10^2$ | $(1.98 \pm 0.17) \times 10^4$ |
| | $K_{\text{i}}$ (mM) | $47.9 \pm 10.0$ | — | $62.5 \pm 21.5$ | — | $49.2 \pm 10.6$ |
| **8a** | $K_{\text{M}}$ (mM) | $0.58 \pm 0.07$ | $0.25 \pm 0.04$ | $0.99 \pm 0.21$ | $1.07 \pm 0.30$ | $0.26 \pm 0.02$ |
| | $k_{\text{cat}}$ ($\text{s}^{-1}$) | $0.16 \pm 0.005$ | $0.23 \pm 0.008$ | $0.16 \pm 0.018$ | $0.04 \pm 0.007$ | $0.65 \pm 0.018$ |
| | $k_{\text{cat}}/K_{\text{M}}$ ($\text{M}^{-1} \text{s}^{-1}$) | $(2.66 \pm 0.26) \times 10^2$ | $(9.26 \pm 1.23) \times 10^2$ | $(1.65 \pm 0.19) \times 10^2$ | $(4.06 \pm 0.60) \times 10^1$ | $(2.47 \pm 0.14) \times 10^3$ |
| | $K_{\text{i}}$ (mM) | — | — | $24.6 \pm 10.9$ | $14.8 \pm 6.7$ | $40.5 \pm 8.5$ |
| **9a** | $K_{\text{M}}$ (mM) | $0.18 \pm 0.03$ | — | — | — | $0.13 \pm 0.02$ |
| | $k_{\text{cat}}$ ($\text{s}^{-1}$) | $0.06 \pm 0.002$ | — | — | — | $0.07 \pm 0.002$ |
| | $k_{\text{cat}}/K_{\text{M}}$ ($\text{M}^{-1} \text{s}^{-1}$) | $(3.41 \pm 0.42) \times 10^2$ | — | — | — | $(5.41 \pm 0.84) \times 10^2$ |
| | $K_{\text{i}}$ (mM) | — | — | — | — | — |
| **10a** | $K_{\text{M}}$ (mM) | $0.10 \pm 0.015$ | $0.17 \pm 0.021$ | $0.04 \pm 0.016$ | $0.62 \pm 0.086$ | $0.06 \pm 0.014$ |
| | $k_{\text{cat}}$ ($\text{s}^{-1}$) | $0.79 \pm 0.03$ | $0.26 \pm 0.001$ | $0.50 \pm 0.04$ | $0.26 \pm 0.016$ | $0.80 \pm 0.04$ |
| | $k_{\text{cat}}/K_{\text{M}}$ ($\text{M}^{-1} \text{s}^{-1}$) | $(8.03 \pm 0.95) \times 10^3$ | $(1.57 \pm 0.15) \times 10^3$ | $(1.27 \pm 0.46) \times 10^4$ | $(4.18 \pm 0.36) \times 10^2$ | $(1.44 \pm 0.30) \times 10^4$ |
| | $K_{\text{i}}$ (mM) | $17.7 \pm 3.5$ | $28.6 \pm 6.9$ | $19.0 \pm 7.7$ | $26.6 \pm 7.8$ | $36.6 \pm 15.1$ |

From the MSMEG_2027 variants, V30Y, V65L, and P122M were selected for kinetic analysis, as these showed improved conversion and *de* for both carvone enantiomers. Mutant enzymes generally outperformed the wildtype in terms of *k*_cat_/*K*_M_. For example, for V65L with **1a** *k*_cat_/*K*_M_ was almost doubled (1.5 × 10^2^ *cf*. 8.2 × 10^1^ M^-1^ s^-1^). V30Y turned over **2a** the most efficiently (*k*cat/*K*M = 1.17 × 10^2^ M^-1^ s^-1^). The V30Y mutation greatly lowered *K*_M_ for all tested substrates, which generally lead to an increase in *k*_cat_/*K*_M_. In several cases, V30Y showed decreased *k*cat, particularly with **7a**, leading to an overall decrease in catalytic efficiency. Moreover, only V30Y showed substrate inhibition with **8a**, **4a** and **10a**.

From the MSMEG_2850 variants, V53I, Q120Y, F121M and Y124M were selected for kinetic analysis, as these variants showed improved conversion and improved or reversed stereoselectivity for both carvone enantiomers. As observed for the MSMEG_2027 variants, the mutant enzymes showed greater *k*_cat_/*K*_M_ than the wildtype. Q120Y increased catalytic activity more than double for **2a** and **1a**, (*k*_cat_/*K*_M_ = 5.2 × 10^1^ M^-1^ s^-1^ and 1.3 × 10^2^ M^-1^ s^-1^, respectively, Table 1). The Y124M mutation reduced *K*_M_ for most of substrates compared to the wildtype. Interestingly, the Y124M mutation increased *k*_cat_/*K*_M_ by almost ten-fold with **8a** (2.5 × 10^3^ M^-1^ s^-1^ *cf*. 2.7 × 10^2^ M^-1^ s^-1^, Table 2), which is greater than that of wildtype MSMEG_2027. The Y124M mutation reduced *K*_M_ (0.26 mM) more than two-fold, and increased *k*_cat_ (0.65 s^-1^) more than fourfold, while acquiring substrate inhibition (*K*_i_ = 40.5 mM). Among the variants only MSMEG_2850-Y124M was able to reduce verbenone **9a** above the detection limit in GC/MS analysis, consistent with the greater catalytic efficiency of MSMEG_2850- Y124M compared to the wildtype.

## Discussion

### Mutations generally deleterious to activity

The conserved serine in the SKGG motif plays an important part in catalysis as shown by MSMEG_2027-S67A having much lower or completely abolished activity with most of the substrates tested. A more striking example of this is that the equivalent S78A mutant of Ddn which showed a ~50% reduction in activity with menadione and complete loss of activity with pretomanid.^17^ The substrates that were turned over appreciably by MSMEG_2027-S67A were those for which the wildtype enzyme has relatively high specific activity.^10^ This was also the case with the M54Q variant and the N11W/Y, F29R, F64N and F121Y variants of MSMEG_2850. Interestingly, the V30R mutation in MSMEG_2027 was not as deleterious as the equivalent mutation in MSMEG_2850 (F29R).

### Selectivity with 1a is predictive of selectivity with other 2-methyl cyclic enone substrates

We compared the effect of each mutation on the selectivity with each substrate. Spearman correlation coefficients showed that substrates could be grouped based upon the effect that mutations had on the stereoselectivity of the reduction. Although there have been many investigations that have examined stereoselectivity across multiple substrates and variants (either mutants or homologues),^27, 45–47^ to our knowledge this is the first use of this statistical approach to quantitatively sequence-function relationships in this way. Remarkably, the substrate groupings were similar between the MSMEG_2027 and MSMEG_2850 variants despite the relatively low sequence identity between these enzymes and the disparate sets of mutations examined. Molecular docking studies indicate that substrates in each set have similar binding orientations to each other and this underlies these correlations. The most notable example is the different carbonyl that acts as the activating group of **7a** with MSMEG_2027 and MSMEG_2850 which mimics the poses of **3a** and **4a**, respectively. Because of this, mutations that affect selectivity with **7a** correlate more strongly with those that affect the substrate that best mimics the binding pose of **7a**. A practical consequence of this correlation is that it suggests that screening of smaller libraries of representative compounds against a panel of enzyme variants may be sufficient to narrow down the number of positions to target *via* saturation mutagenesis or rational design.

### Identification of selectivity and reactivity hot spots

We identified several regions that have a dramatic effect on the activity and/or the selectivity of the enzymes. Aromatic substitution of V30 in MSMEG_2027 resulted in reversed stereoselectivity with **8a**, particularly with tryptophan. In addition, the Y120A/Q and Y126M variants decreased or reversed the stereoselectivity with **1a** and **2a** while slightly increased selectivity with **8a**. In contrast, Y124M and F121M decreased or switched stereoselectivity with carvones (**1a** and **2a**) and 2-methyl substituted cyclic enones (**3a** and **5a**). MSMEG_2850-V53I showed decreased selectivity with **8a** (37% *de*) while showing improved selectivity with 2-methyl substituted cyclic enones **1a**, **2a**, **3a** and **5a**. F121Y retained 95% conversion with **8a** but had no detectable activity with **1a** and **2a**.

### Effect of mutations in different genetic backgrounds

Several of the mutations we examined resulted in residues in MSMEG_2027 being mutated to the equivalent position in the MSMEG_2850 or vice versa, providing insight into the effect of these residues in different genetic backgrounds. One such example is MSMEG_2850-F121Y that remarkably showed low or no detectable conversion of most substrates despite this mutation corresponding to Y123 in MSMEG_2027. Both Phe and Tyr are conserved at this position (Figure 1) and the equivalent mutation in Ddn (Y133F) retains much of the wildtype activity with menadione and nitroimidazoles.^18^ Hydrogen bonding between tyrosine and the pyrimidine ring of F_420_ likely increases the redox potential of the cofactor, thereby leading to lower turnover as is seen with the T37A mutant of OYE2.^48^ This is consistent with the F121M variant retaining or exceeding the activity of the wildtype with most substrates. This hydrogen bonding may be compensated for by other interactions in enzymes that poses a tyrosine at this position, such as MSMEG_2027, Ddn and FDR-Rh2. Future studies could examine the effect of the reciprocal Tyr→Phe mutation in these enzymes.

Another set of reciprocal variants is MSMEG_2027-Y120A and MSMEG_2850- A118Y. The Y120A variant generally resulted in lower conversions compared to wildtype MSMEG_2027, whereas the A118Y variant generally showed higher conversions than wildtype MSMEG_2850. Interestingly, both mutations resulted in diminished stereoselectivity with most substrates, illustrating a complex interplay between selectivity and activity. This is further shown with the MSMEG_2027-Y120Q variant generally showing loss of activity and selectivity compared to wildtype MSMEG_2027, whereas MSMEG_2850-A118Q showed comparable or even greater activity and selectivity than the wildtype. The V30F mutation in MSMEG_2027 converts this position to the equivalent residue in MSMEG_2850 (F29) and resulted in reversed selectivity with **8a**.

Disparate effects of the same residue in different genetic backgrounds have also been seen in the OYE family. For example, the W116I mutation in *Saccharomyces pastorianus* OYE 1 resulted in opposite stereochemical outcomes for several substrates, whereas the close homologue OYE 2.6 from *Pichia stipitis* that has Ile at this position has identical selectivity to the wildtype OYE 1.^49^ Similarly, wildtype TsER and DrER produce (*S*)-**4b**, whereas YqjM produces (*R*)-**4b** despite having identical active site residues to these homologues.^33^ The NCR T25G/W66T mutant retains the wildtype preference for (2*R*)-**2b**, whereas the equivalent double mutant flips selectivity in RmER, TsER and DrER to (2*S*)-**2b**. Strong epistatic effects have been noted in protein engineering efforts previously and highlight the limitations of rational protein engineering approaches.^50^

### Potential effects on protein stability and dimerization

The active forms of both MSMEG_2027 and MSMEG_2850 are monomeric, however both enzymes form head-to-head dimers in the absence of cofactor that partially occlude the active site.^10, 51^ As a consequence, many of the residues targeted for mutagenesis here are involved in the dimer interface as well as substrate binding. It is unclear what effect these mutations may have on dimer formation and protein stability. Perturbation of dimer formation is of interest for MSMEG_2027 given that it forms the basis of a recently reported chemically dependent protein de-dimerization tool for studying protein-protein interactions.^51^ The OYE homologue 12-oxophytodienoate reductase 3 (OPR3) from tomato forms a self-inhibiting dimer through insertion of a loop into the active site that is similar to what is observed in MSMEG_2027 and MSMEG_2850.^52^ Interestingly two point mutations were identified that completely abolished OPR3 dimer formation, although the protomers do not undergo the large conformational change seen in MSMEG_2027 upon dissociation.^52^

## Conclusion

Using a structure-based rational design approach we constructed a targeted library of 46 single point mutations across two wildtype sequences for improved activity with (*R*)-carvone reduction. We identified several variants with improved activity and selectivity using this substrate, along with some variants that unexpectedly showed reversed stereoselectivity. We then screened this library against a panel of nine additional substrates containing prochiral alkenes and determined the effect of each mutation on the conversion and stereoselectivity with each substrate. Using Spearman correlation, we found that substrates could be grouped together based on which mutations affected stereoselectivity. Notably, there was a negative correlation between α-methyl cyclic enones and (*R*)-myrtenal. We found that the correlation between ketoisophorone and α-methyl cyclic enones differs between the two genetic backgrounds and that this reflects a preference for which carbonyl serves as the activating group between these enzymes.

We identified several regions where mutations can have a dramatic effect on activity. These include V30 and Y126 of MSMEG_2027 as well as V53 and Y124 of MSMEG_2850. Mutation of the conserved tyrosine to methionine reversed selectivity with α-methyl cyclic enones, while mutation of V30 to aromatic residues reversed selectivity with (*R*)-myrtenal. Computational docking suggested that this was most likely due to altered facial selectivity of protonation following hydride transfer, rather than from the substrate adopting a “flipped” binding mode. We hypothesize that this Tyr→Met mutation disrupts the hydrophobic shield created by three conserved aromatic residues in the active site and that this facilitates entry of solvent molecules that protonate the substrate. The Y124M mutation also enabled MSMEG_2850 to convert verbenone above the detection limit of the assay.

Overall, our strategy produced several variants with improved activity and increased or reversed selectivity with many of the substrates. It also yielded useful information regarding the determinants of stereoselectivity (that appear to differ from the OYE family) that may guide future engineering efforts. The OYE family follow a bi–bi pingpong mechanism, and therefore need to facilitate NADPH oxidation as well as substrate reduction.^45^ FDORs do not share this requirement as their cofactor is reduced externally and may therefore be more amenable to engineering without this constraint.

## Acknowledgements

This work was supported by the CSIRO SynBio FSP, the ARC Centre of Excellence in Synthetic Biology (CE200100029), and the ARC Centre of Excellence in Peptide and Protein Science (CE200100012). JA was supported by the Australian Government Research Training Program Scholarship (RTP). We thank Adam Carroll, Joe Boileau, Thy Truong, and Anitha Jeyasingham of the Australian National University RSC/RSB Joint Mass Spectrometry Facility for assistance with GC/MS analyses.

