## Supplementary Information for "Active site mutations of F_420_-dependent alkene reductases reverse stereoselectivity"

### Investigating the mechanism and specificity of $F_{420}$-dependent oxidoreductases through mutagenesis

[1] Research School of Chemistry, Australian National University, Canberra, ACT 2601, Australia
[2] Natural Products Research Center, Korea Institute of Science and Technology (KIST), Gangneung 25451, Republic of Korea
[3] School of Biology and Environmental Science, Queensland University of Technology, Brisbane, QLD 4000, Australia
[4] ARC Centre of Excellence in Synthetic Biology, Queensland University of Technology, Brisbane, Queensland 4000, Australia
[5] School of Chemistry, Monash University, Melbourne, Victoria 3800, Australia
[6] Environment, Commonwealth Scientific and Industrial Research Organisation, Clayton, Victoria 3168, Australia.
[7] CSIRO Advanced Engineering Biology Future Science Platform, GPO Box 1700, Canberra, ACT 2601, Australia
[8] ARC Centre of Excellence in Synthetic Biology, Research School of Chemistry, Australian National University, Canberra, ACT 2601, Australia
[9] ARC Centre of Excellence for Innovations in Peptide & Protein Science, Research School of Chemistry, Australian National University, Canberra, ACT 2601, Australia

#### Materials and methods

##### Chemicals

The following chemicals were obtained from Sigma–Aldrich (St. Louis, MO): (*R*)-(−)-carvone, **1a**; (*S*)-(+)-carvone, **2a**; (+)-dihydrocarvone, **1b**; 2-methyl-2-cyclohexen-1-one, (**3a**), (1*R*)-(−)-myrtenal (**8a**), (+)-*cis*-myrtanol, (1*S*)-(−)-verbenone (**9a**), citral (**10a**), (*S*)-(−)-citronellal ((*S*)-**10b**), (*R*)-(+)-citronellal ((*R*)-**10b**), geraniol and nerol. Ketoisophorone (**7a**) was purchased from AK Scientific (Union City, CA); 3-methyl-2-cyclohexen-1-one (**4a**), 3-methylcyclohexanone (**4b**), 2-methylcyclohexanone (**3b**), 3-methyl-2-cyclopenten-1-one (**6a**), 3-methylcyclopentanone (**6b**), 2-methyl-2-cyclopenten-1-one (**5a**), and 2-methylcyclopentanone (**5b**) were obtained from Enamine (Kiev, Ukraine). (−)-*trans*-myrtanol was purchased from abcr GmbH (Karlsruhe, Germany). (+)-*cis*-myrtanal (*cis*-**8b**), (+)-*trans*-myrtanal (*trans*-**8b**) geranial (*trans*-**10a**) and neral (*cis*-**10a**) were prepared from the corresponding alcohols with Dess–Martin periodinane as previously described.[1] Cofactor $F_{420}$ was prepared using *E. coli* BL21(DE3) and *M. smegmatis* mc$^2$ 4517 overexpressing $F_{420}$ biosynthetic genes as described previously.[2, 3] Other chemicals and solvents were sourced from commercial suppliers as analytical or molecular biology grade and used without further purification unless otherwise stated.

##### Cloning, expression, and purification of enzymes

Plasmids for expression of wild type MSMEG_2027 (UniProtKB accession #A0QU01) and MSMEG_2850 (UniProtKB accession #A0QW82), and $F_{420}$-dependent glucose-6-phosphate dehydrogenase (Fgd) were as previously described.[1] Using these sequences as templates, mutants were designed and ordered from Twist Bioscience (CA, USA) cloned into pET28a(+) between the *Nco*I and *Xho*I sites. The His6-tag and TEV cleavage sites of pETMCSIII were retained in these constructs. *E. coli* BL21(DE3) chemically competent cells were prepared using the Mix & Go kit (Zymo Research) following the manufacturers' instructions or were purchased from New England Biolabs. All enzymes were expressed in modified auto-induction media as previously described using 50 µg mL$^{-1}$ kanamycin or 100 µg mL$^{-1}$ ampicillin as appropriate.[1, 4] Proteins were purified as previously described with the following modifications that facilitated the parallel purification of all variants. Cells were resuspended in 500 µl lysis buffer containing 50 mM sodium phosphate, pH 8.0, 300 mM NaCl, 25 mM imidazole, and 1× BugBuster® protein extraction reagent (Merck). Following 30 minutes of incubation at room temperature the cell suspension was centrifuged at 13 000*g* for 10 min, and the supernatant was loaded onto a HisPur Ni-NTA 96-well Spin Plates (Thermo scientific). Samples were washed following the manufacturer's recommendations and eluted with buffer containing 250 mM imidazole. The eluted proteins were buffer-exchanged into storage buffer (50 mM Tris-HCl, 300 mM NaCl, 10% glycerol, pH 8.0) using Amicon® Ultracentrifugation filters (10 kDa, Merck Millipore). SDS-PAGE confirmed soluble expression of the mutants (Figure S3). Flash frozen aliquots of purified protein were stored at −80 °C until use.

**GC/MS analysis**

Enzyme reactions were performed in Eppendorf tubes for 18 hours containing 50 μM $F_{420}$, 3 μM FDOR, 1.2 mM substrate, 8.3 mM G6P and 1.4 μM Fgd in assay buffer (50 mM Tris, 300 mM NaCl, pH 8.0) in a final volume of 30 μL. Reactions were extracted with an equal volume of ethyl acetate containing 200 μM mesitylene as an internal standard. Samples were analyzed using GC/MS (7890A GC system, 5975C MSD, Agilent, Santa Clara, CA). Columns, injection programs, and retention times are described in Table S9. Substrate and products were identified using the NIST MS spectrum library and the identification was confirmed with standard compounds or with previously reported retention indices.[5-8]

**Computational methods**

MSMEG_2027 and MSMEG_2850 were modelled using previously determined X-ray structures (PDB 6WTA and 8D4W, respectively). The $F_{420}$-4 molecule of the MSMEG_2027:$F_{420}$-4 structure was integrated into MSMEG_2850 structure, followed by Prime loop refinement (Schrodinger LLC, NY). Both structures were parameterized with the OPLS4 forcefield and subjected to energy minimization using Desmond (Schrodinger LLC, NY) with the default settings.[9] Following minimization, waters were removed from the structures, and substrates parameterized with OPLS4 were docked into the active sites using the induced fit protocol in Maestro 2022-01 release (Schrodinger LLC, NY).

Residue Scanning Calculation in Schrödinger-biologics was used to predict changes in binding affinity, stability, and prime energy (Schrödinger LLC, NY). PROPKA and Epik were used to predict the ionization states of proteins and ligands, respectively. Optimizing hydrogen-bonding networks was done with ProtAssign. For mutant variants, the mutational site was switched into an appropriate residue and subjected to energy minimization using Desmond. The binding free energy ($\Delta G_{bind}$)

between enzymes and substrates was calculated using prime MM-GBSA with implicit VSGB solvation model, OPLS4 force field, and default Prime settings.

Apoenzyme structures of three other previously characterized FDOR-A1s, FDR-Mha (UniProtKB accession #K5BI92), FDR-Rh1 (UniProtKB accession #Q0S7M3), and FDR-Rh2 (UniProtKB accession #Q0S5L4), were obtained from the AlphaFold2 database.[10] They were combined with $F_{420}$ and prepared as described above. The T-coffee web server[11] was used to produce multiple sequence alignments, and the ESPript web server[12] was used to render them. The sap_pair[13] and clustalw_msa[14] methods were used for structural alignments using the Expresso method in T-coffee.[15]

**Determination of enzyme kinetics**

$F_{420}$ was reduced enzymatically with 0.2–1.0 µM Fgd and excess G6P in assay buffer that had been degassed under nitrogen. Fgd was removed using Amicon Ultra centrifuge filters (10 kDa, Merck Millipore) and the resulting solution of $F_{420}H_2$ was used immediately. Stock solutions (200 mM) of each substrate was prepared in methanol, except **7a** which was dissolved in the assay buffer. Enzyme was preincubated with $F_{420}H_2$ for five minutes before assays were initiated by addition of substrate. 100 µL reactions contained 0.1 µM FDOR, 10 µM $F_{420}H_2$ and 0.078–10 mM substrate. Activity was measured fluorometrically by following the reoxidation of $F_{420}H_2$ ($\lambda_{ex}$ = 420 nm, $\lambda_{em}$ = 470 nm) on an Infinite 200 PRO plate reader and i-control software (Tecan Trading AG, Switzerland) using black 96-well plates (Costar, Corning, NY).[16] Assays were conducted in at technical duplicate. Prism v.9.3 (GraphPad Software Inc., La Jolla, CA) was used to determine apparent steady-state kinetic parameters based on the standard Michaelis–Menten equation (1) or Michaelis–Menten with substrate inhibition equation (2).

$$v = \frac{k_{\mathrm{cat}}[E][S]}{K_{\mathrm{M}} + [S]} \quad (1)$$

$$v = \frac{k_{\mathrm{cat}}[E][S]}{K_{\mathrm{M}} + [S]\left(1 + \frac{[S]}{K_{\mathrm{i}}}\right)} \quad (2)$$

**Residual activity and selectivity over long term storage**

Loss of activity during long term storage is a frequent problem with biocatalysts. We compared activity and stereoselectivity of enzymes stored at 4 °C for 3 months with that of freshly prepared enzymes (Figure S16). Proteolytic cleavage was observed with most variants by SDS-PAGE, whereas the wild-type enzymes remained intact (Figure S17). Despite this, considerable activity remained after three months. The most significant attenuation of activity and selectivity was seen with 3-methyl substituted substrates **4a** and **6a**, for which the baseline activity is low. Interestingly V30F showed a substantial drop of *ee* (38.0%) against **8a**, while unexpectedly V53I yielded remarkable increase of *ee* (191.3% increase to a total *ee* value of 62.2%), after 3-month-storage. Overall, although less stable than wild type, the mutant enzymes are relatively stable at 4 °C for 3 months.

#### Supplementary tables

**Table S1.** MM-GBSA energy parameters for MSMEG_2027 variants with **1a**

| Mutant | ΔAffinity[a] | ΔStability[a] | ΔPrime Energy[a] |
|---|---|---|---|
| V30M | −5.34 | −5.26 | −12.03 |
| V30R | −2.88 | −3.45 | −37.11 |
| V30I | −2.46 | −0.03 | 6.92 |
| V30W | −2.34 | 1.01 | −13.81 |
| V30F | −2.31 | 2.29 | −1.55 |
| L31Y | −2.23 | 1.18 | −12.9 |
| V30Y | −2.09 | 3.06 | −9.49 |
| L31M | −1.88 | −5.31 | −10.94 |
| L31F | −1.81 | 1.22 | −4.33 |
| Y126M | −1.1 | −4.63 | 4.97 |
| L31R | −0.76 | −3.75 | −36.06 |
| V65L | −0.01 | −5.25 | −4.13 |
| P122M | 0.04 | −7.04 | −38.95 |
| P122R | 3.21 | −6.2 | −62.79 |
| W11N | 5.49 | 13.99 | −16.53 |
| V12N | 0.17 | 16.31 | −31.46 |
| M54Q | 1.45 | 18.95 | −23.63 |
| Y120Q | 5.96 | 28.64 | −5.82 |
| Y120A | 6.87 | 38.31 | 46.85 |
| S67A | 3.86 | 3.14 | 4.52 |

[a] $kcal\ mol^{-1}$

**Table S2.** MM-GBSA energy parameters for MSMEG_2850 variants with **1a**

| Mutant | ΔAffinity[a] | ΔStability[a] | ΔPrime Energy[a] |
|---|---|---|---|
| V53M | −8.23 | −13.82 | −19.76 |
| F29R | −6.01 | −6.51 | −34.96 |
| M10Y | −5.11 | 0.43 | −5.75 |
| E30M | −4.31 | −0.83 | 20.94 |
| Q120M | −4.21 | −5.8 | 33.74 |
| Q120R | −4.15 | −4.04 | 9.16 |
| V53I | −4.1 | −9.5 | −2.02 |
| Q120L | −3.79 | −2.11 | 42.27 |
| V53Y | −3.7 | −3.28 | −15.68 |
| A118F | −3.58 | 29.43 | 26.54 |
| V53L | −3.49 | −4.22 | −4.56 |
| M10R | −3.48 | −7.45 | −29.71 |
| N11W | −3.33 | −3.32 | 30.69 |
| A118M | −2.86 | −9.78 | −14.39 |
| N11Y | −2.85 | 0.66 | 36.27 |
| A118L | −2.76 | 33.57 | 36.32 |
| A118Y | −2.62 | 58.15 | 48.42 |
| A118H | −2.36 | 14.55 | 3.9 |
| A118R | −2.22 | 16.76 | −12 |
| E30W | −2.12 | 0.53 | 13.7 |
| F121M | −1.94 | −1.27 | −2.62 |
| F121Y | −1.5 | 2.89 | −5.73 |
| Y124M | −0.76 | −7.73 | 1.65 |
| A118Q | −0.46 | −0.01 | −44.03 |
| F64N | −0.03 | 20.62 | −22.15 |
| Q120Y | 0.96 | 0.17 | 36.33 |

[a] kcal $mol^{-1}$

**Table S3.** Screening of prochiral substrates with MSMEG_2027 variants

| Variant | Substrate | | | | | | | | | | | |
|---|---|---|---|---|---|---|---|---|---|---|---|---|
| | **1a** | **2a** | **3a** | **4a** | **5a** | **6a** | **7a** | **8a** | **9a** | **10a** | *cis*-**10a** | *trans*-**10a** |
| | c (%)[a] | c (%)[a] | c (%)[a] | c (%)[a] | c (%)[a] | c (%)[a] | c (%)[a] | c (%)[a] | c (%)[a] | c (%)[a] | c (%)[a] | c (%)[a] |
| | *de* (%)[b] | *de* (%)[b] | *ee* (%)[c] | *ee* (%)[c] | *ee* (%)[c] | *ee* (%)[c] | *ee* (%)[c] | *de* (%)[b] | *ee* (%)[c] | *ee* (%)[c] | *ee* (%)[c] | *ee* (%)[c] |
| WT | 89.1 | 88.4 | >99 | 30.9 | 70.8 | 3.6 | >99 | >99 | ND[d] | 76.2 | 95.0 | 95.4 |
| | 85.3 | 91.8 | 88.0 | >99 | 60.9 | >99 | 82.7 | 63.6 | ND[d] | >99 | >99 | >99 |
| W11N | 73.5 | 83.2 | 22.4 | 2.3 | 8.5 | ND[d] | >99 | 95.7 | ND[d] | 31.5 | — | — |
| | 54.8 | 87.1 | 69.9 | >99 | 49.3 | ND[d] | 89.3 | 72.0 | ND[d] | >99 | — | — |
| V12N | 91.9 | 90.2 | >99 | 32.7 | 83.0 | 4.5 | >99 | >99 | ND[d] | 81.5 | — | — |
| | 83.4 | 90.8 | 89.6 | >99 | 63.0 | >99 | 82.1 | 49.3 | ND[d] | >99 | — | — |
| V30M | 86.9 | 88.8 | >99 | 10.0 | 81.7 | ND[d] | >99 | >99 | ND[d] | 49.8 | — | — |
| | 84.0 | 92.4 | 87.6 | >99 | 80.2 | ND[d] | 83.7 | 58.6 | ND[d] | >99 | — | — |
| V30R | 34.8 | 48.2 | 65.4 | 2.7 | 30.0 | ND[d] | 98.2 | >99 | ND[d] | 9.2 | — | — |
| | 76.0 | 87.4 | 86.0 | >99 | 60.9 | ND[d] | 88.8 | 57.3 | ND[d] | >99 | — | — |
| V30I | 87.8 | 86.5 | >99 | 40.0 | 81.2 | 3.5 | >99 | >99 | ND[d] | 87.2 | — | — |
| | 76.3 | 91.4 | 87.6 | >99 | 57.1 | >99 | 87.2 | 44.5 | ND[d] | >99 | — | — |
| V30W | 89.9 | 92.4 | >99 | 27.6 | 85.3 | ND[d] | 67.3 | >99 | ND[d] | 82.0 | 95.0 | 91.3 |
| | 83.6 | 89.7 | 89.4 | >99 | 58.7 | ND[d] | 84.4 | −41.1 | ND[d] | >99 | >99 | >99 |
| V30F | 90.7 | 91.9 | >99 | 88.3 | >99 | 5.7 | 80.8 | >99 | ND[d] | 84.0 | 97.7 | 91.8 |
| | 92.9 | 91.3 | 87.7 | >99 | 42.1 | >99 | 84.7 | −21.5 | ND[d] | >99 | >99 | >99 |
| V30Y | 90.3 | 92.9 | >99 | 89.5 | >99 | 7.0 | 81.9 | >99 | ND[d] | 82.4 | 96.4 | 91.8 |
| | 87.3 | 92.3 | 89.7 | >99 | 45.6 | >99 | 87.9 | −16.9 | ND[d] | >99 | >99 | >99 |
| L31Y | 91.7 | 89.6 | >99 | 21.3 | 75.5 | 3.3 | 98.9 | >99 | ND[d] | 85.2 | — | — |
| | 83.5 | 91.7 | 86.8 | >99 | 58.4 | >99 | 82.6 | 46.2 | ND[d] | >99 | — | — |
| L31M | 91.7 | 91.0 | >99 | 26.8 | 81.7 | 2.7 | >99 | >99 | ND[d] | 78.0 | — | — |
| | 77.3 | 91.1 | 88.3 | >99 | 57.1 | >99 | 86.7 | 55.7 | ND[d] | >99 | — | — |
| L31F | 90.5 | 90.5 | >99 | 25.1 | 80.1 | 3.0 | >99 | >99 | ND[d] | 87.4 | — | — |
| | 84.0 | 89.4 | 88.8 | >99 | 57.6 | >99 | 86.9 | 53.6 | ND[d] | >99 | — | — |
| L31R | 85.8 | 90.3 | >99 | 18.9 | 76.7 | 3.9 | >99 | >99 | ND[d] | 84.8 | — | — |
| | 81.4 | 91.2 | 86.8 | >99 | 65.1 | >99 | 88.0 | 49.7 | ND[d] | >99 | — | — |
| M54Q | ND[d] | 1.3 | 6.2 | 2.2 | 5.0 | ND[d] | 21.8 | 43.8 | ND[d] | 6.5 | — | — |
| | ND[d] | 48.5 | 29.2 | >99 | 32.9 | ND[d] | 79.0 | 56.6 | ND[d] | >99 | — | — |
| V65L | 94.3 | 92.6 | >99 | 32.2 | 95.4 | 4.0 | >99 | >99 | ND[d] | 86.2 | — | — |
| | 89.2 | 92.3 | 79.9 | >99 | 78.8 | >99 | 87.4 | 33.3 | ND[d] | >99 | — | — |
| S67A | ND[d] | 1.3 | 7.8 | 1.1 | 3.7 | ND[d] | 62.0 | 75.4 | ND[d] | 12.0 | — | — |
| | ND[d] | 31.1 | 34.4 | >99 | 33.8 | ND[d] | 61.8 | 85.7 | ND[d] | >99 | — | — |
| Y120Q | 19.3 | 22.4 | 41.6 | 11.0 | 24.9 | ND[d] | 98.0 | >99 | ND[d] | 59.4 | — | — |
| | 36.6 | 57.3 | 75.2 | >99 | 56.4 | ND[d] | 89.0 | 76.2 | ND[d] | >99 | — | — |
| Y120A | 69.3 | 53.0 | 97.0 | 5.2 | 84.7 | ND[d] | >99 | >99 | ND[d] | 57.7 | — | — |
| | 55.0 | 45.9 | 81.9 | >99 | 86.7 | ND[d] | 74.3 | 81.2 | ND[d] | >99 | — | — |
| P122M | 91.0 | 91.8 | >99 | 28.4 | 91.1 | 3.9 | >99 | >99 | ND[d] | 83.2 | 92.6 | 96.0 |
| | 91.8 | 91.7 | 90.9 | >99 | 74.6 | >99 | 89.5 | 48.1 | ND[d] | >99 | >99 | >99 |
| P122R | 90.8 | 90.7 | >99 | 25.0 | 84.9 | 4.7 | >99 | >99 | ND[d] | 86.1 | — | — |
| | 81.2 | 91.9 | 85.1 | >99 | 70.5 | >99 | 89.7 | 33.4 | ND[d] | >99 | — | — |
| Y126M | 57.0 | 83.1 | >99 | 12.9 | 70.8 | 2.6 | >99 | >99 | ND[d] | 77.8 | — | — |
| | −17.3 | 28.0 | −13.6 | >99 | −18.5 | >99 | 54.4 | 76.2 | ND[d] | >99 | — | — |

[a] conversion [b] diastereomeric excess [c] enantiomeric excess [d] ND not determined, conversion below detection limit. Stereoselectivity is reported with the major product of the wild type enzyme assigned a positive value, and the alternative product assigned a negative value.

**Table S4.** Screening of prochiral substrates with MSMEG_2850 variants

| | Substrate | | | | | | | | | | | |
|---|---|---|---|---|---|---|---|---|---|---|---|---|
| Variant | **1a** | **2a** | **3a** | **4a** | **5a** | **6a** | **7a** | **8a** | **9a** | **10a** | *cis*-**10a** | *trans*-**10a** |
| | c (%)[a] *de* (%)[b] | c (%)[a] *de* (%)[b] | c (%)[a] *ee* (%)[c] | c (%)[a] *ee* (%)[c] | c (%)[a] *ee* (%)[c] | c (%)[a] *ee* (%)[c] | c (%)[a] *ee* (%)[c] | c (%)[a] *de* (%)[b] | c (%)[a] *ee* (%)[c] | c (%)[a] *ee* (%)[c] | c (%)[a] *ee* (%)[c] | c (%)[a] *ee* (%)[c] |
| WT | 84.2 | 64.0 | 62.6 | >99 | 62.6 | 85.9 | >99 | >99 | ND[d] | 93.7 | 99.0 | 98.9 |
| | 60.0 | 83.0 | 62.9 | >99 | 44.5 | >99 | 87.0 | 72.8 | ND[d] | 25.0 | >99 | −52.3 |
| M10Y | 66.2 | 55.8 | 27.9 | >99 | 41.0 | 39.8 | >99 | 96.9 | ND[d] | 93.8 | — | — |
| | 50.3 | 82.5 | 64.0 | >99 | 40.2 | >99 | 86.0 | 79.6 | ND[d] | 22.3 | — | — |
| M10R | 21.8 | 22.1 | 11.7 | 92.5 | 12.5 | 6.9 | >99 | 90.1 | ND[d] | 92.3 | 97.2 | 98.3 |
| | 46.5 | 82.8 | 41.8 | >99 | 36.4 | >99 | 87.5 | 79.1 | ND[d] | −17.1 | >99 | −70.7 |
| N11W | ND[d] | ND[d] | 7.4 | 4.0 | ND[d] | ND[d] | >99 | 15.5 | ND[d] | 46.6 | — | — |
| | ND[d] | ND[d] | 28.9 | >99 | ND[d] | ND[d] | 89.6 | 62.8 | ND[d] | 14.5 | — | — |
| N11Y | ND[d] | ND[d] | 6.5 | 3.8 | ND[d] | ND[d] | >99 | 14.1 | ND[d] | 66.5 | 44.8 | 83.4 |
| | ND[d] | ND[d] | 40.1 | >99 | ND[d] | ND[d] | 89.3 | 65.9 | ND[d] | −24.1 | 83.8 | −66.8 |
| F29R | 4.7 | 15.5 | 10.8 | 73.7 | 5.9 | 3.5 | >99 | 79.6 | ND[d] | 86.3 | 97.4 | 94.2 |
| | >99 | 78.7 | 43.6 | >99 | 51.3 | >99 | 88.0 | 66.6 | ND[d] | 61.3 | >99 | 26.6 |
| E30M | 82.9 | 56.1 | 51.5 | >99 | 57.5 | 69.3 | >99 | 97.3 | ND[d] | 91.9 | — | — |
| | 62.9 | 79.5 | 68.6 | >99 | 41.1 | >99 | 87.9 | 79.2 | ND[d] | 22.2 | — | — |
| E30W | 72.9 | 47.0 | 36.6 | >99 | 42.2 | 51.2 | >99 | >99 | ND[d] | 90.1 | — | — |
| | 61.8 | 76.7 | 64.2 | >99 | 42.7 | >99 | 90.0 | 77.6 | ND[d] | 34.2 | — | — |
| V53M | 81.6 | 63.2 | 73.1 | >99 | 60.7 | 84.6 | >99 | >99 | ND[d] | 95.3 | — | — |
| | 57.2 | 82.5 | 64.3 | >99 | 38.2 | >99 | 86.0 | 77.5 | ND[d] | 37.8 | — | — |
| V53I | 93.0 | 76.1 | 93.0 | >99 | 93.7 | 17.3 | >99 | >99 | ND[d] | 95.9 | 98.7 | 97.5 |
| | 80.4 | 87.0 | 85.0 | >99 | 79.4 | >99 | 85.1 | 37.0 | ND[d] | 77.5 | >99 | 67.1 |
| V53Y | 81.2 | 58.0 | 50.0 | >99 | 62.5 | 44.5 | >99 | >99 | ND[d] | 97.8 | — | — |
| | 71.4 | 77.7 | 74.7 | >99 | 42.7 | >99 | 89.4 | 60.4 | ND[d] | 38.9 | — | — |
| V53L | 84.3 | 62.8 | 66.3 | >99 | 71.8 | 74.3 | >99 | >99 | ND[d] | 93.6 | — | — |
| | 64.4 | 79.5 | 71.4 | >99 | 46.3 | >99 | 89.9 | 71.4 | ND[d] | 39.3 | — | — |
| F64N | 7.6 | 2.3 | 6.0 | 67.3 | 3.8 | 1.8 | >99 | 58.0 | ND[d] | 94.4 | — | — |
| | 38.8 | 26.4 | 41.9 | >99 | 49.9 | >99 | 91.0 | 71.4 | ND[d] | 35.0 | — | — |
| A118F | 90.8 | 75.8 | 90.2 | >99 | 89.8 | 85.3 | >99 | >99 | ND[d] | 92.3 | — | — |
| | 41.2 | 63.4 | 67.8 | >99 | 48.9 | >99 | 89.1 | 71.2 | ND[d] | 14.6 | — | — |
| A118M | 88.7 | 73.4 | 83.8 | >99 | 90.7 | 82.1 | >99 | >99 | ND[d] | 92.7 | — | — |
| | 60.6 | 80.8 | 62.7 | >99 | 41.1 | >99 | 86.8 | 70.8 | ND[d] | 27.4 | — | — |
| A118L | 88.3 | 71.3 | 76.8 | >99 | 90.2 | 47.9 | >99 | >99 | ND[d] | 96.1 | — | — |
| | 71.5 | 78.1 | 68.8 | >99 | 48.1 | >99 | 89.7 | 71.0 | ND[d] | 18.5 | — | — |
| A118Y | 90.0 | 74.2 | 88.5 | >99 | 92.2 | 88.1 | >99 | 94.8 | ND[d] | 95.9 | — | — |
| | 26.5 | 60.4 | 61.7 | >99 | 50.0 | >99 | 87.9 | 61.2 | ND[d] | 17.6 | — | — |
| A118H | 89.1 | 66.5 | 72.9 | >99 | 72.3 | 55.8 | >99 | >99 | ND[d] | 95.6 | — | — |
| | 50.4 | 76.0 | 63.3 | >99 | 46.4 | >99 | 88.4 | 76.8 | ND[d] | 8.0 | — | — |
| A118R | 87.4 | 67.7 | 72.2 | >99 | 76.2 | 61.1 | >99 | >99 | ND[d] | 94.7 | — | — |
| | 68.2 | 82.3 | 67.0 | >99 | 52.3 | >99 | 89.1 | 69.5 | ND[d] | 6.3 | — | — |
| A118Q | 89.6 | 72.0 | 83.8 | >99 | 81.3 | 84.4 | >99 | >99 | ND[d] | 95.2 | — | — |
| | 60.6 | 80.1 | 69.6 | >99 | 45.9 | >99 | 90.2 | 80.3 | ND[d] | 23.7 | — | — |
| Q120M | 75.2 | 51.4 | 58.5 | >99 | 63.8 | 47.0 | >99 | >99 | ND[d] | 94.7 | — | — |
| | 45.2 | 77.0 | 57.4 | >99 | 38.5 | >99 | 89.8 | 84.6 | ND[d] | 13.1 | — | — |
| Q120R | 81.2 | 60.4 | 59.8 | >99 | 57.4 | 54.2 | >99 | >99 | ND[d] | 94.8 | — | — |
| | 63.8 | 78.4 | 64.8 | >99 | 47.0 | >99 | 89.1 | 72.4 | ND[d] | 18.6 | — | — |
| Q120L | 89.6 | 75.8 | 84.4 | >99 | 91.4 | 95.8 | >99 | >99 | ND[d] | 94.6 | 99.4 | 99.1 |
| | 43.0 | 80.7 | 66.9 | >99 | 49.6 | >99 | 91.5 | 83.3 | ND[d] | −3.0 | >99 | −74.9 |
| Q120Y | 95.0 | 87.2 | 97.2 | >99 | 97.5 | 97.5 | >99 | 97.0 | ND[d] | 95.0 | 99.4 | 98.2 |
| | 68.1 | 86.0 | 77.2 | >99 | 40.3 | >99 | 91.7 | 74.7 | ND[d] | 64.3 | >99 | 49.9 |
| F121M | 86.6 | 80.5 | 69.5 | 92.8 | 77.0 | 3.7 | >99 | 93.2 | ND[d] | 95.3 | — | — |
| | −18.3 | −14.2 | 37.0 | >99 | 60.0 | >99 | 91.1 | 85.0 | ND[d] | 6.3 | — | — |
| F121Y | ND[d] | ND[d] | 4.8 | 53.3 | ND[d] | 1.7 | >99 | 95.0 | ND[d] | 94.7 | — | — |
| | ND[d] | ND[d] | 21.7 | >99 | ND[d] | >99 | 91.4 | 84.4 | ND[d] | 15.1 | — | — |
| Y124M | 81.4 | 58.3 | 57.2 | >99 | 75.4 | 36.6 | >99 | >99 | 10.0 | 95.9 | — | — |
| | −36.2 | 18.1 | −5.1 | >99 | 17.8 | >99 | 91.7 | 86.9 | >99 | 0.5 | — | — |

[a]conversion [b]diastereomeric excess [c]enantiomeric excess [d]ND not determined, conversion below detection limit. Stereoselectivity is reported with the major product of the wild type enzyme assigned a positive value, and the alternative product assigned a negative value.

**Table S5.** Spearman correlation coefficients between substrates for MSMEG_2027 variants

| | **1a** | **2a** | **3a** | **4a**[#] | **5a** | **6a**[#] | **7a** | **8a** | **9a**[#] | **10a**[#] |
|---|---|---|---|---|---|---|---|---|---|---|
| **1a** | 1.00 | 0.72*** | 0.66** | — | 0.22 | — | 0.02 | −0.65*** | — | — |
| **2a** | | 1.00 | 0.53* | — | 0.48* | — | 0.39 | −0.63*** | — | — |
| **3a** | | | 1.00 | — | 0.30 | — | 0.21 | −0.57** | — | — |
| **4a** | | | | — | — | — | — | — | — | — |
| **5a** | | | | | 1.00 | — | 0.24 | −0.13 | — | — |
| **6a** | | | | | | — | — | — | — | — |
| **7a** | | | | | | | 1.00 | −0.34 | — | — |
| **8a** | | | | | | | | 1.00 | — | — |
| **9a** | | | | | | | | | — | — |
| **10a** | | | | | | | | | | — |

* $p < 0.05$ ** $p < 0.01$ *** $p < 0.005$ [#]not calculated, no variation

**Table S6.** Spearman correlation coefficients between substrates for MSMEG_2850 variants

| | **1a** | **2a** | **3a** | **4a**# | **5a** | **6a**# | **7a** | **8a** | **9a**# | **10a** |
|---|---|---|---|---|---|---|---|---|---|---|
| **1a** | 1.00 | 0.49* | 0.66*** | — | 0.17 | — | −0.23 | −0.57*** | — | 0.61*** |
| **2a** | | 1.00 | 0.43* | — | −0.21 | — | −0.48* | −0.14 | — | 0.34 |
| **3a** | | | 1.00 | — | 0.13 | — | −0.13 | −0.27 | — | 0.53*** |
| **4a** | | | | — | — | — | — | — | — | — |
| **5a** | | | | | 1.00 | — | 0.05 | −0.49* | — | 0.05 |
| **6a** | | | | | | — | — | — | — | — |
| **7a** | | | | | | | 1.00 | 0.42* | — | −0.21 |
| **8a** | | | | | | | | 1.00 | — | −0.35 |
| **9a** | | | | | | | | | — | — |
| **10a** | | | | | | | | | | 1.00 |

* $p < 0.05$ ** $p < 0.01$ *** $p < 0.005$ #not calculated, no variation

**Table S7.** GC/GS methods and retention times

| Method | Substrates | Column[a] | Conditions | $R_t$ (min) |
|---|---|---|---|---|
| 1 | **1a** | 1 | 90 °C for 3 min; 1.5 °C min$^{-1}$ until 123 °C; hold for 5 min; 10 °C min$^{-1}$ until 180 °C; hold for 2 min | **1a**, 24.38<br>*cis*-1**b**, 19.69<br>*trans*-1**b**, 20.57 |
| | **8a** | | | **8a**, 20.01<br>*cis*-8**b**, 20.25<br>*trans*-8**b**, 21.58 |
| | **9a** | | | **9a**, 17.92<br>**9b**, 18.73 |
| 2 | **2a** | 2 | 90 °C for 3 min; 1.5 °C min$^{-1}$ until 123 °C; hold for 5 min; 10 °C min$^{-1}$ until 180 °C; hold for 2 min | **2a**, 24.47<br>*trans*-**2b**, 17.51<br>*cis*-**2b**, 18.49 |
| 3 | **3a** | 3 | 80 °C for 6.5 min; 10 °C min$^{-1}$ until 130 °C; hold for 3 min | **3a**, 10.68<br>(*S*)-**3b**, 10.04<br>(*R*)-**3b**, 10.15 |
| 4 | **4a** | 1 | 40 °C for 0 min; 5 °C min$^{-1}$ until 95 °C; hold for 10 min; 10 °C min$^{-1}$ until 150 °C; hold for 10 min | **4a**, 24.25<br>(*R*)-**4b**, 15.76<br>(*S*)-**4b**, 16.07 |
| 5 | **5a** | 1 | 40 °C for 0 min; 5 °C min$^{-1}$ until 68 °C; hold for 15 min; 10 °C min$^{-1}$ until 160 °C; hold for 2 mins | **5a**, 17.28<br>(*R*)-**5b**, 12.32<br>(S)-**5b**, 12.95 |
| 6 | **6a** | 1 | 40 °C for 0 min; 2 °C min$^{-1}$ until 55 °C; hold for 40 min; 10 °C min$^{-1}$ until 150 °C; hold for 5 mins | **6a**, 33.17<br>(*R*)-**6b**, 23.27<br>(*S*)-**6b,** 23.90 |
| 7 | **7a** | 3 | 90 °C for 2 min; 4 °C min$^{-1}$ until 115 °C; hold for 10 min; 10 °C min$^{-1}$ until 180 °C; hold for 2 min | **7a**, 15.62<br>(*R*)-**7b**, 18.05<br>(*S*)-**7b**, 18.75 |
| 8 | **10a** | 1 | 70 °C for 2 min; 1 °C min$^{-1}$ until 90 °C; hold for 10 min; 4 °C min$^{-1}$ until 130 °C; hold for 2 mins; 10 °C min$^{-1}$ until 150 °C; hold for 1 min | *cis*-**10a**, 41.40<br>*trans*-**10a**, 43.77<br>(*S*)-**10b**, 32.65<br>(*R*)-**10b**, 32.82 |

[a] Column 1: CycloSil-B (30 m × 0.25 mm × 0.25 µm); Column 2: DB-WAX column (30 m × 0.25 mm × 0.5 µm); Column 3: CP-Chirasil-Dex CB (25 m × 0.32 mm × 0.25 µm).

#### Supplementary figures

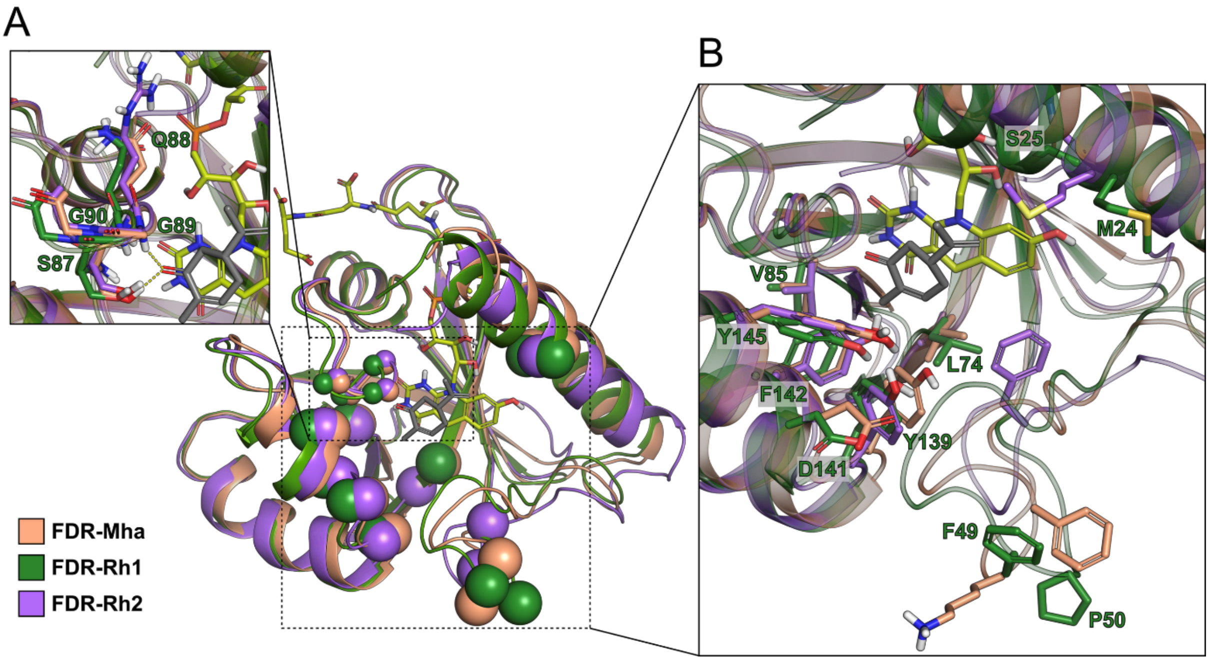

**Figure S1**. Holoenzyme models of FDR-Mha, FDR-Rh1, and FDR-Rh2

Structures were prepared using AlphaFold models and energy minimization as per Materials and Methods. For clarity, only the $F_{420}$-4 molecule and **1a** computationally integrated into of FDR-Mha are shown. **1a** and $F_{420}$ appear in grey and yellow green, respectively. (A) The side chains of S(Q/R)GG motif forming oxyanion hole with 1a are shown as sticks. Hydrogen bonds are represented by yellow dashed lines. (B) The side chains corresponding to the mutational targets of MSMEG_2027 and MSMEG_2850 are shown. Residues of FDR-Rh1 are labelled.

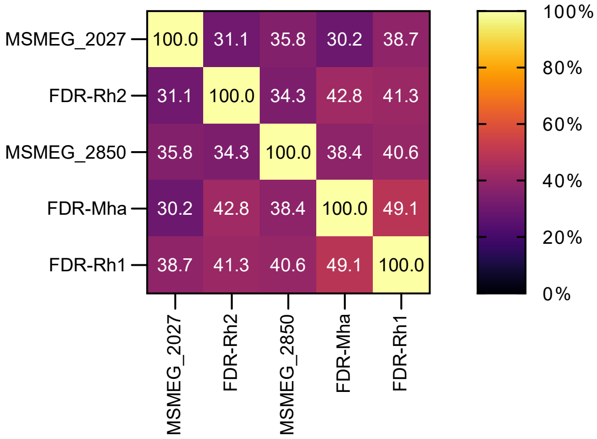

**Figure S2.** Pairwise sequence identify matrix of FDOR-A1 enzymes

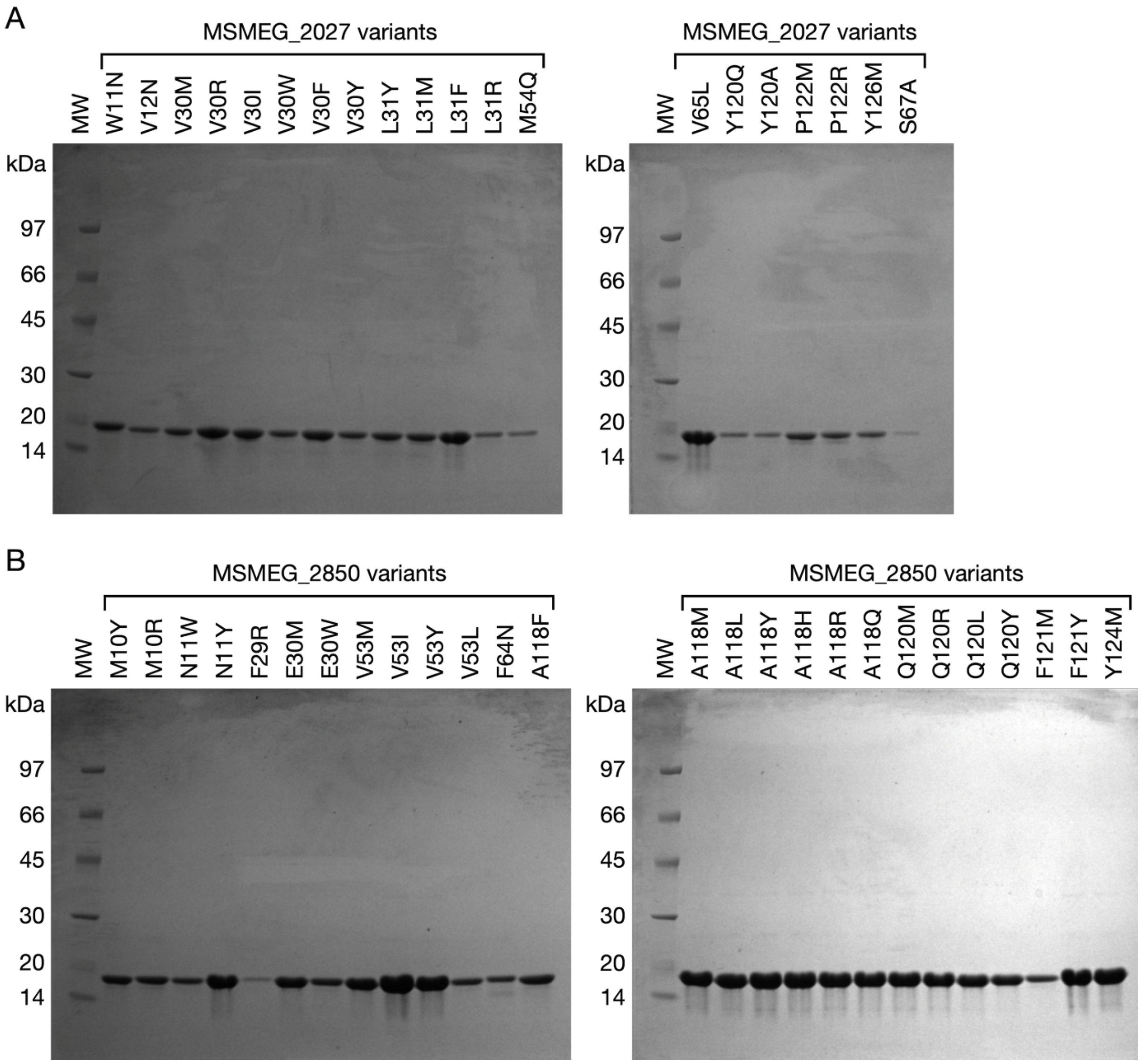

**Figure S3**. SDS–PAGE of purified recombinantly expressed variants

Variants of MSMEG_2027 (A) and MSMEG_2850 (B) were recombinantly expressed and purified using Ni-NTA affinity chromatography. MW, Amersham low molecular weight marker (GE Healthcare).

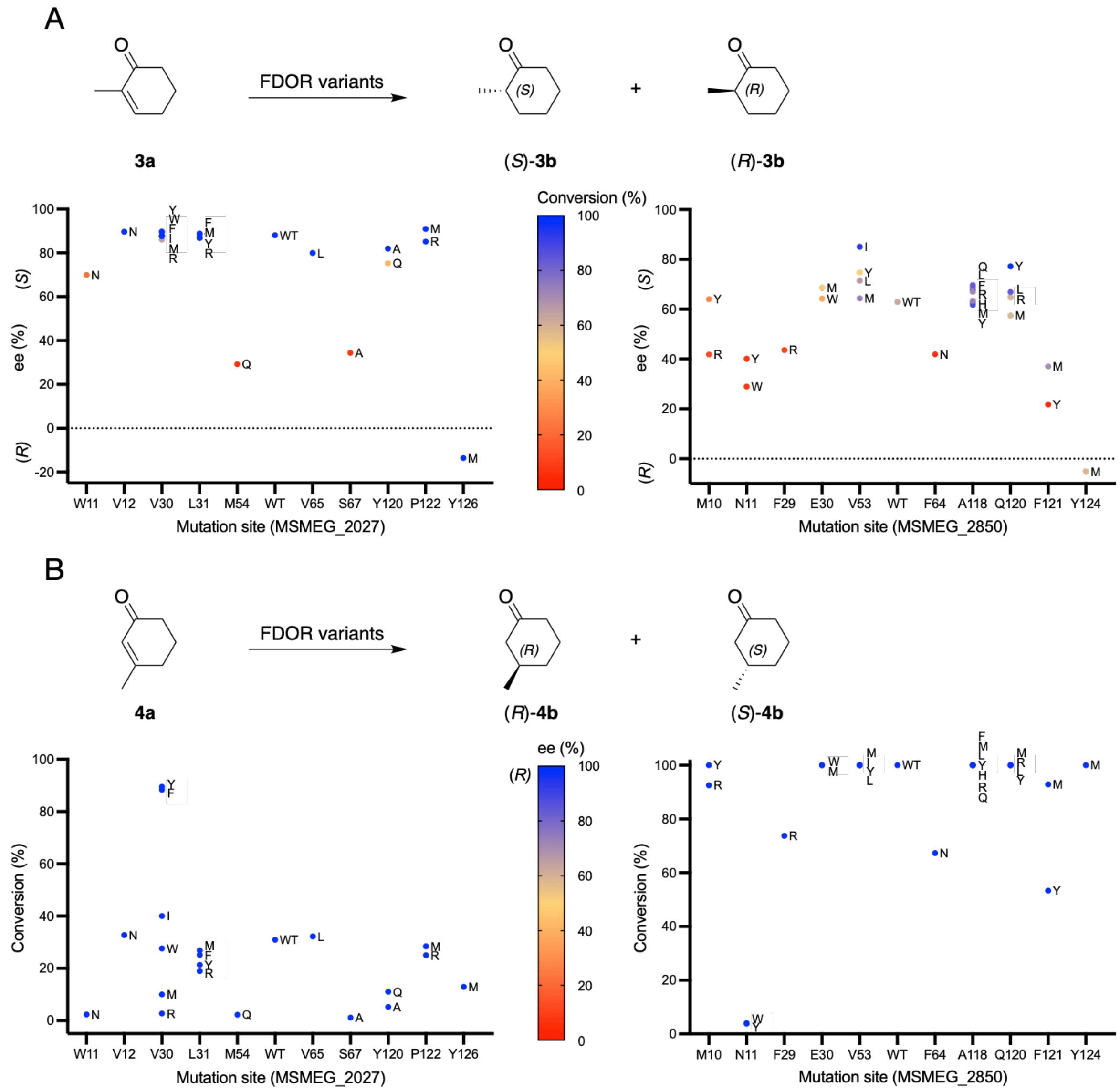

**Figure S4.** Variant screen with cyclohexeneone substrates **3a** and **4a**

Conversions and selectivity of variants of MSMEG_2027 and MSMEG_2850 with (A) 2-methyl-cyclohexenone **3a** and (B) 3-methyl-cyclohexenone **4a**.

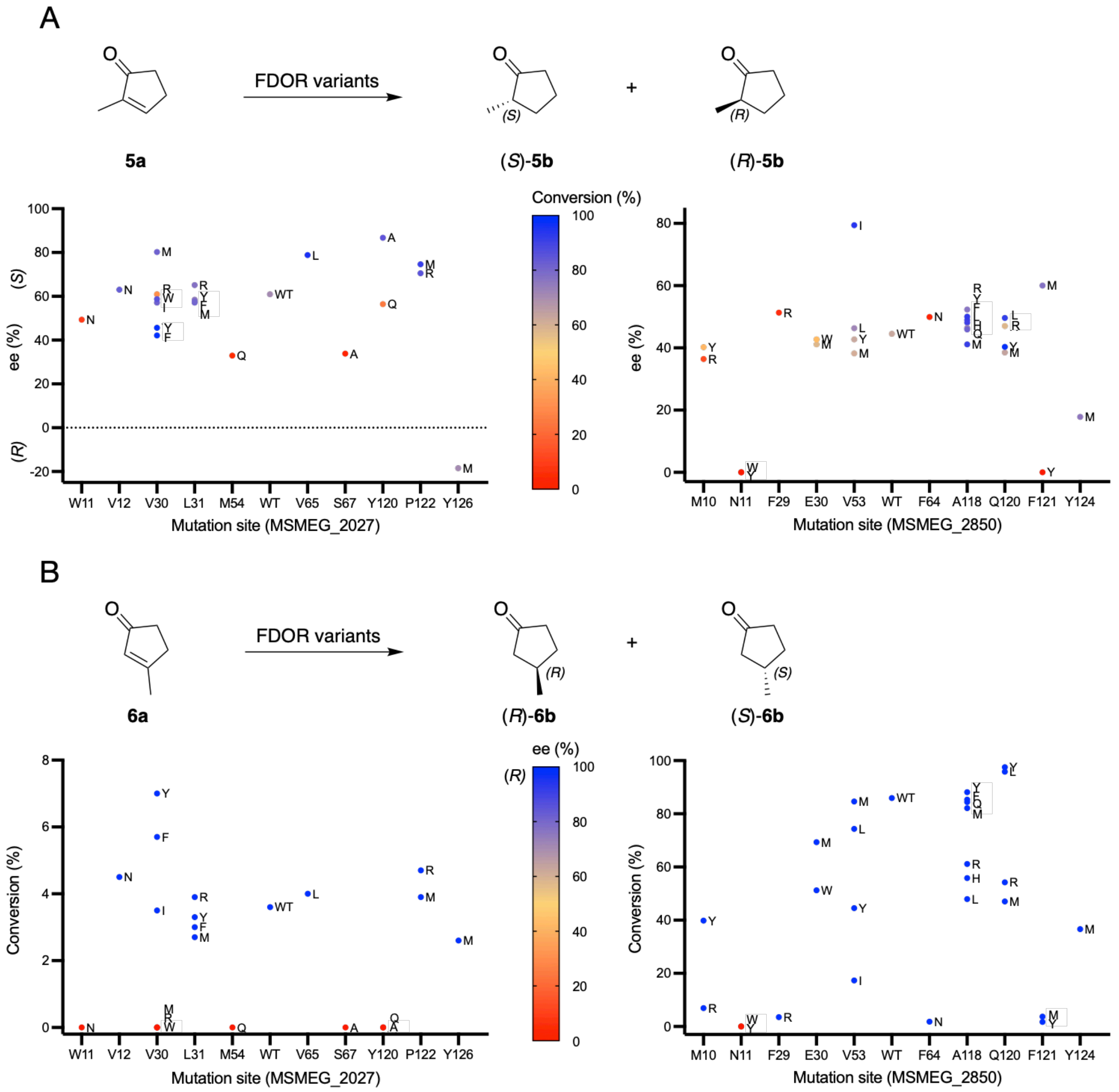

**Figure S5.** Variant screen with cyclopenteneones **5a** and **6a**

Conversions and selectivity of variants of MSMEG_2027 and MSMEG_2850 with (A) 2-methyl-cyclopentenone **5a** and (B) 3-methyl-cyclopentenone **5a**.

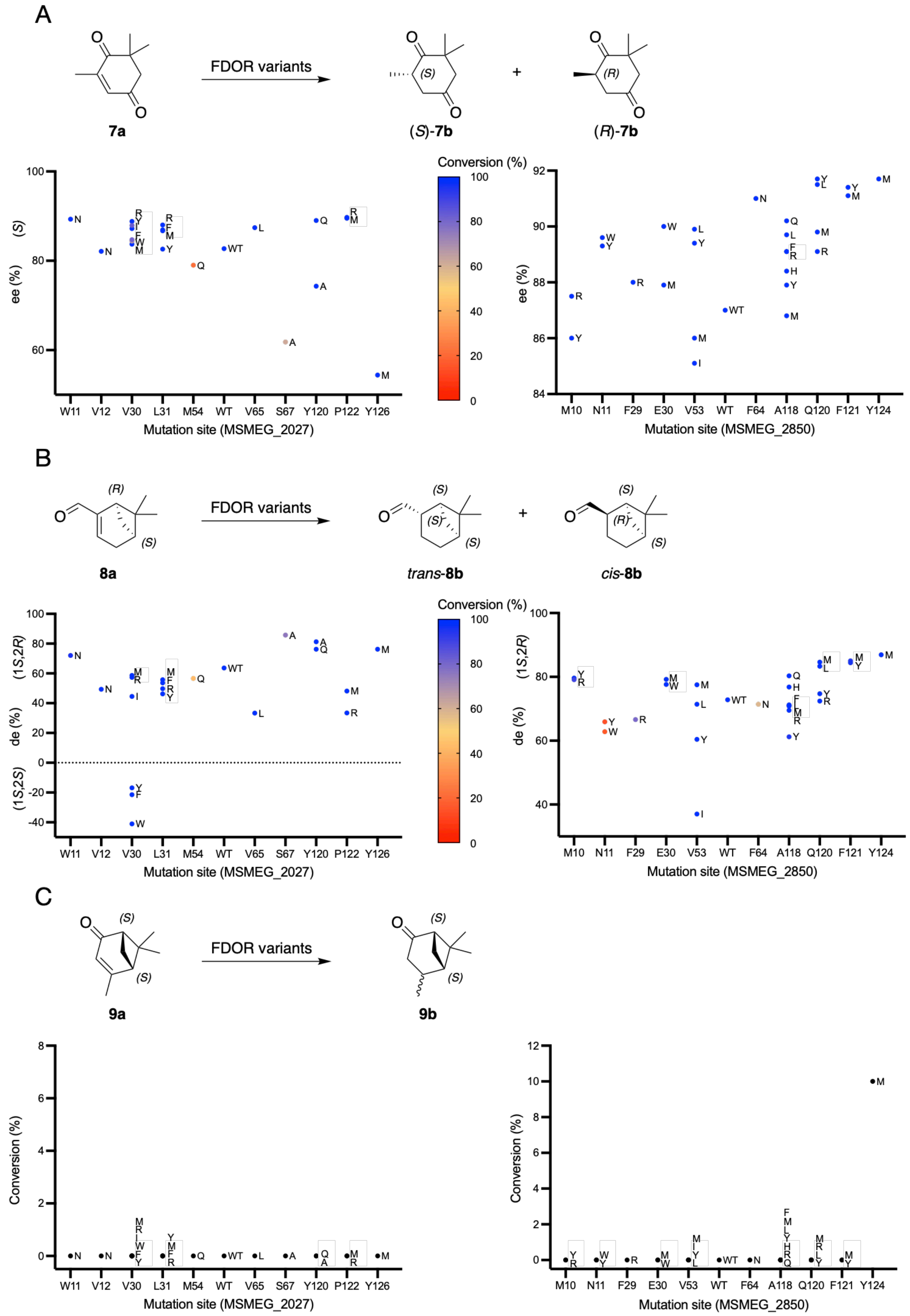

**Figure S6.** Variant screen with substrates **7a, 8a**, and **9a**

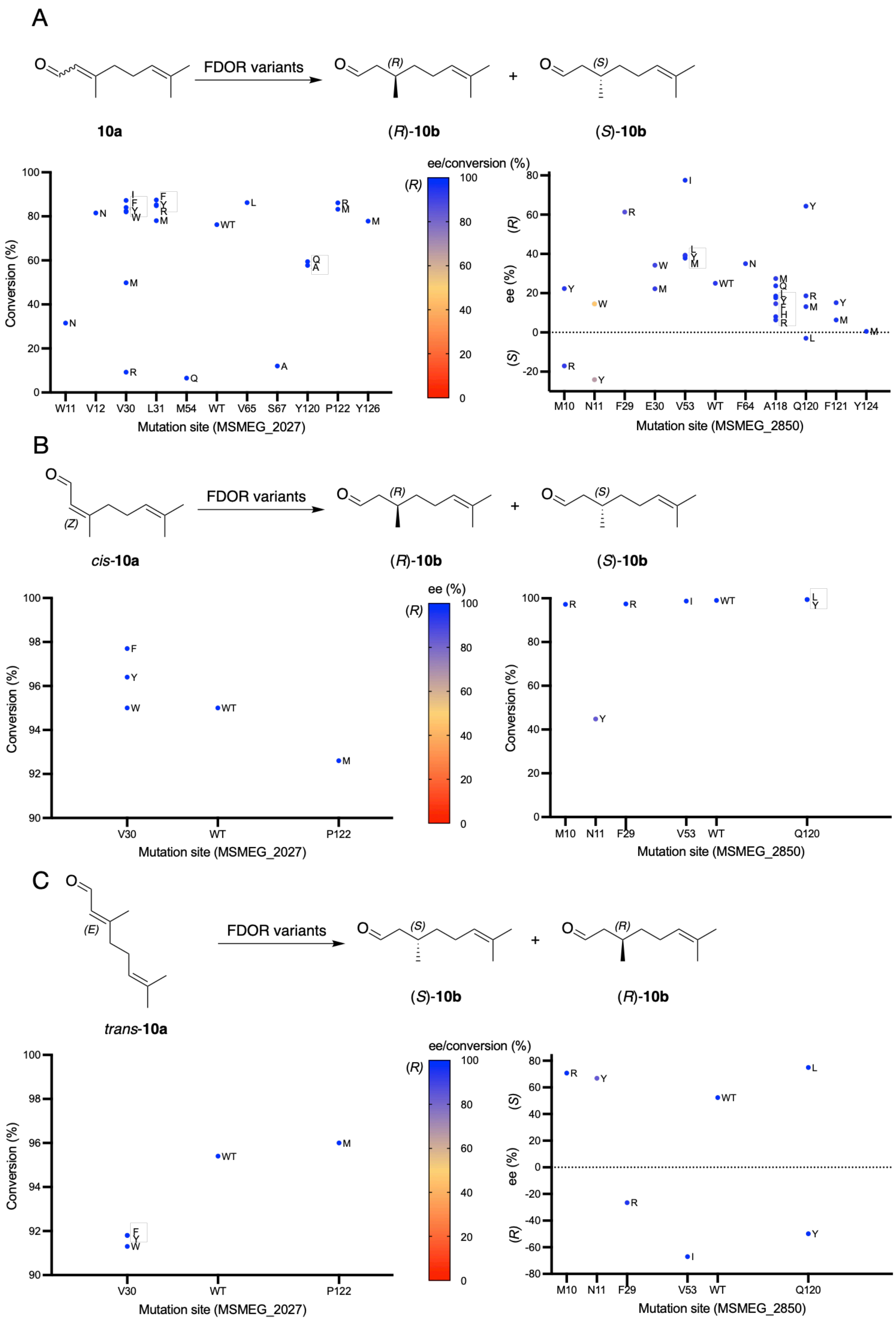

**Figure S7.** Variant screen with commercial **10a** and its *cis* and *trans* isomers

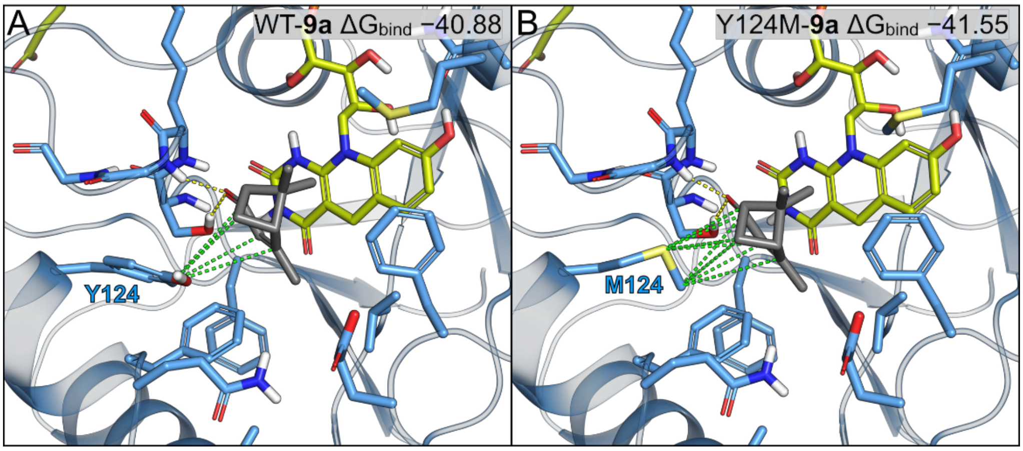

**Figure S8.** Docking of **9a** and with WT and the Y124M variant of MSMEG_2850

Hydrogen bonds are represented by yellow dashed lines, and relevant hydrophobic interactions by green dashed lines. MM-GBSA scores for each pose are shown ($\Delta G_{bind}$, kcal mol$^{-1}$).

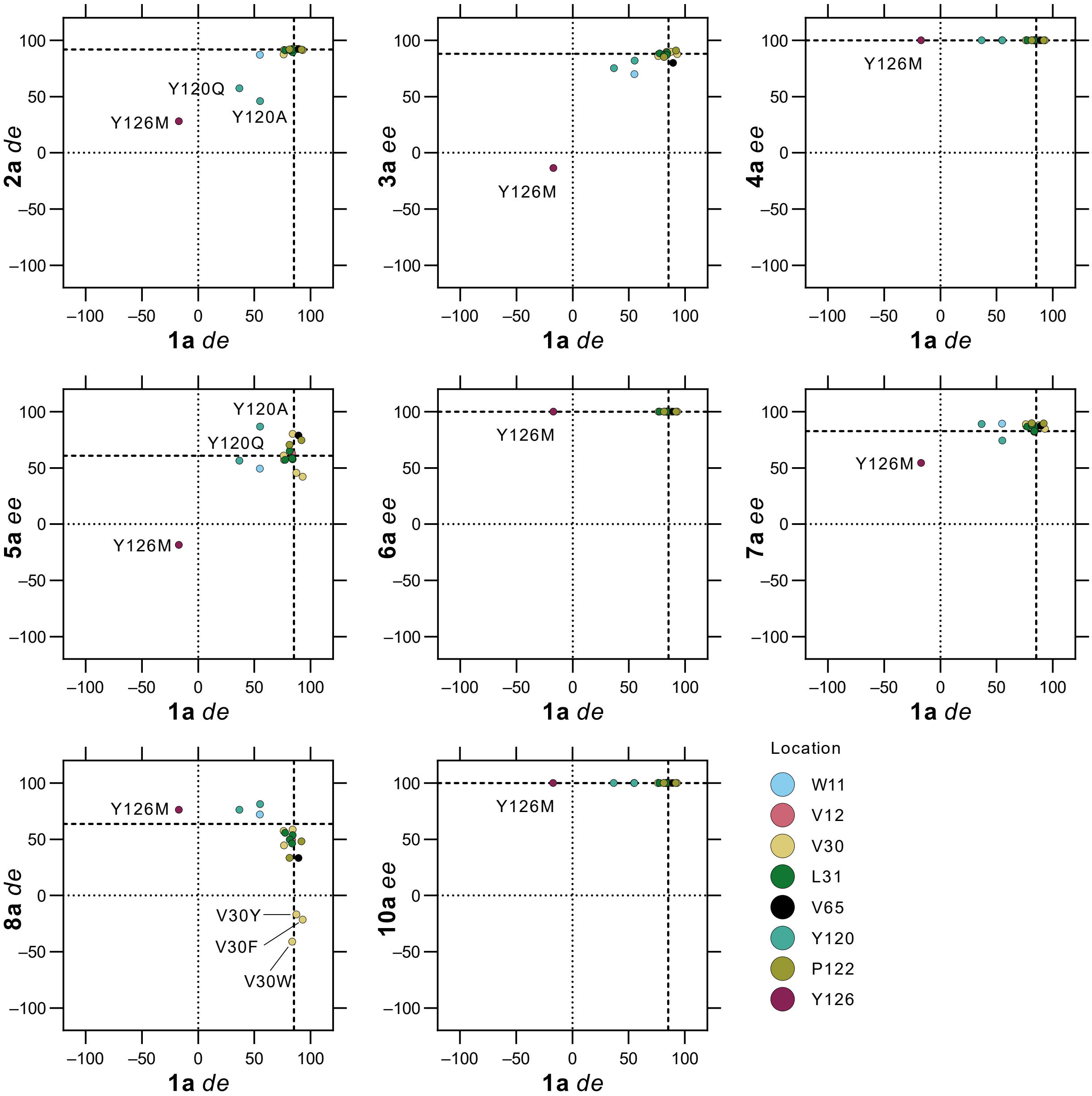

**Figure S9.** Pairwise selectivity comparisons with MSMEG_2027 variants

Values for wild type are indicated with dashed lines. Variants are colored by their position and selected variants are labelled.

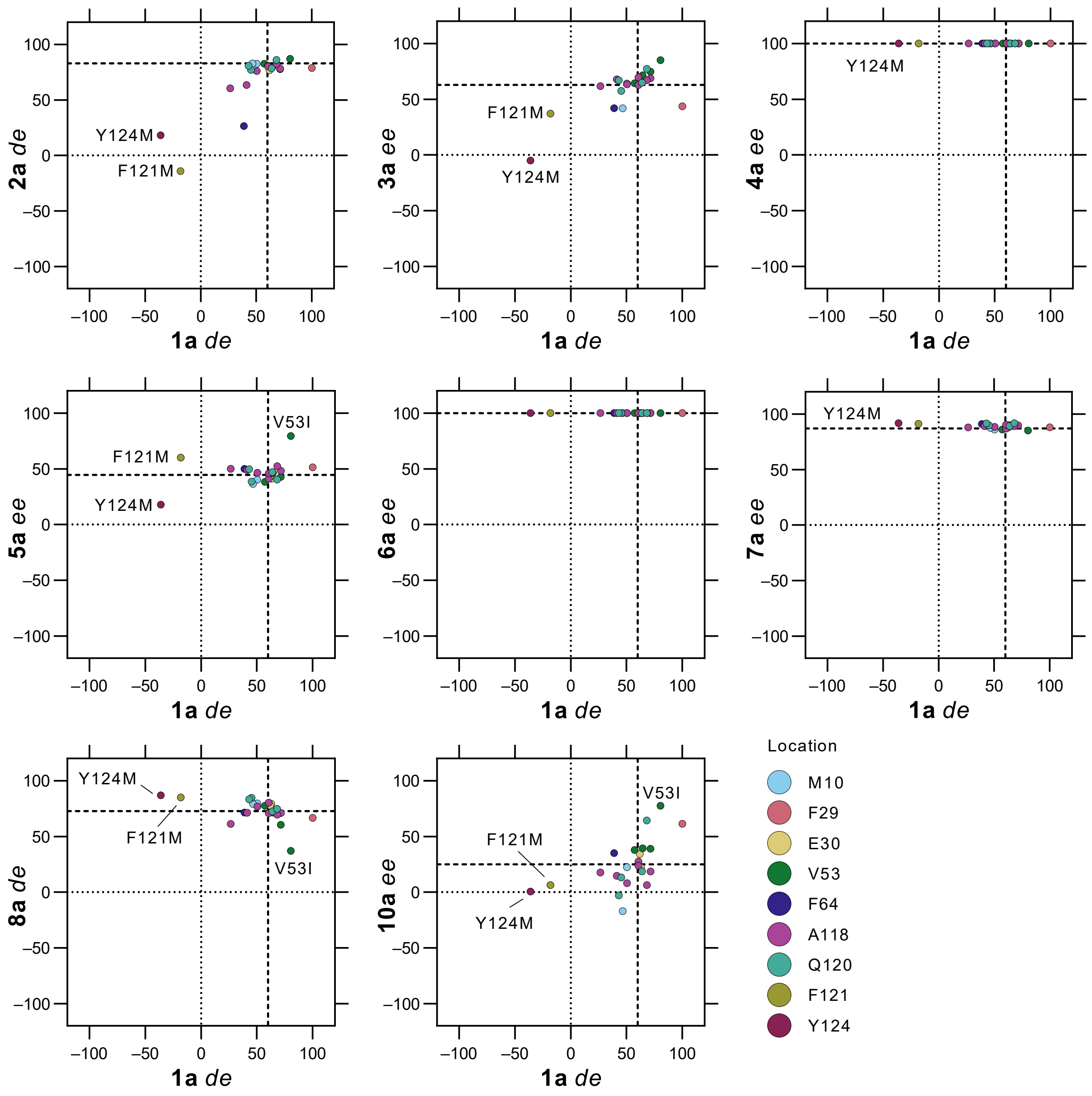

**Figure S10.** Pairwise selectivity comparisons with MSMEG_2850 variants

Values for wild type are indicated with dashed lines. Variants are colored by their position and selected variants are labelled.

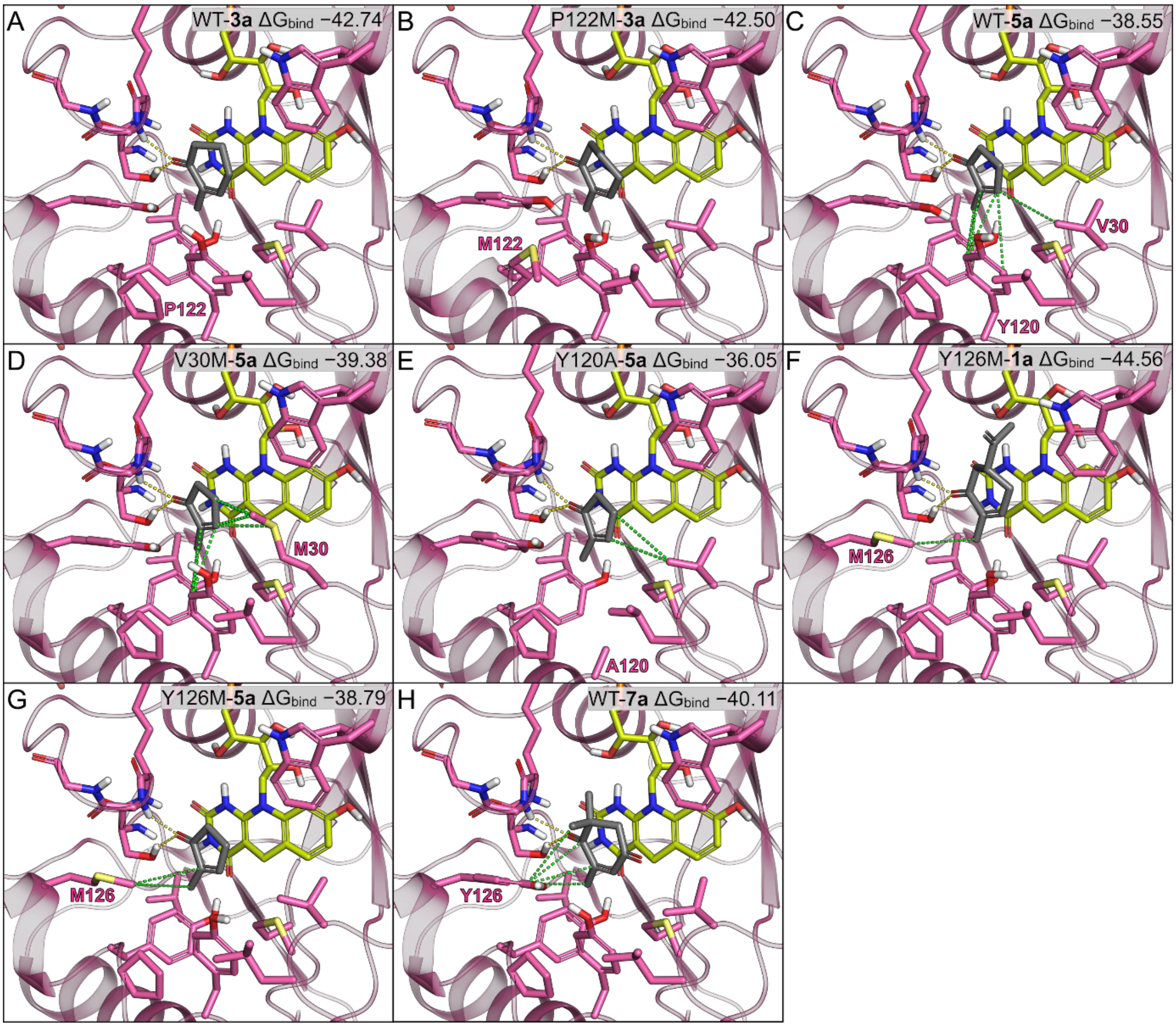

**Figure S11.** Docking poses of MSMEG_2027 variants with **1a**, **3a**, **5a** and **7a**

MSMEG_2027 variants were docked using the induced fit protocol in Schrodinger. Hydrogen bonds are represented by yellow dashed lines, and relevant hydrophobic interactions by green dashed lines. Binding affinities ($\Delta G_{bind}$, kcal mol$^{-1}$) were calculated using MM-GBSA.

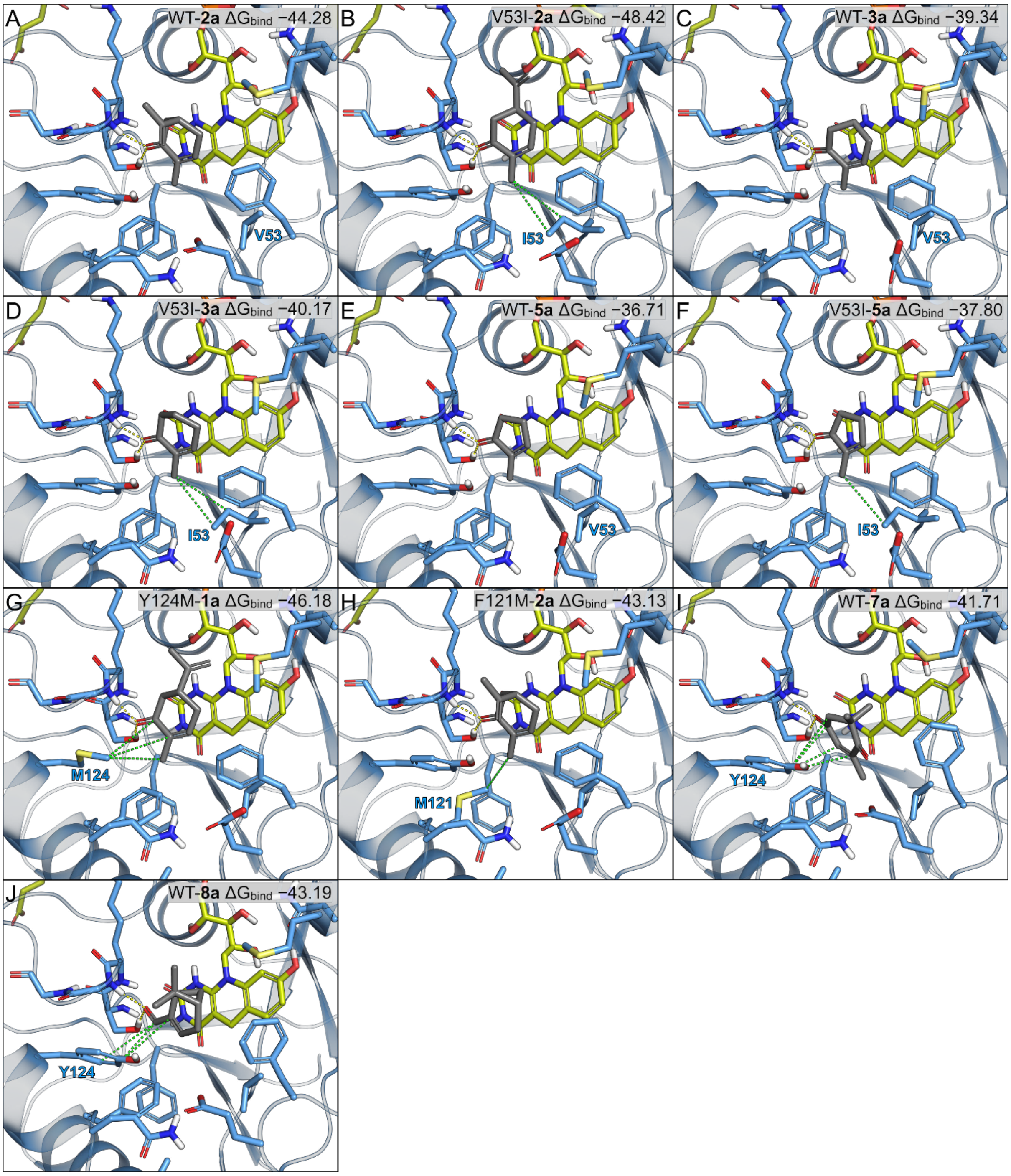

**Figure S12.** Docking poses of MSMEG_2850 variants with **1a**, **2a**, **3a**, **5a**, **7a** and **8a** MSMEG_2850 variants were docked using the induced fit protocol in Schrodinger. Hydrogen bonds are represented by yellow dashed lines, and relevant hydrophobic interactions by green dashed lines. Binding affinities ($\Delta G_{bind}$, kcal mol$^{-1}$) were calculated using MM-GBSA.

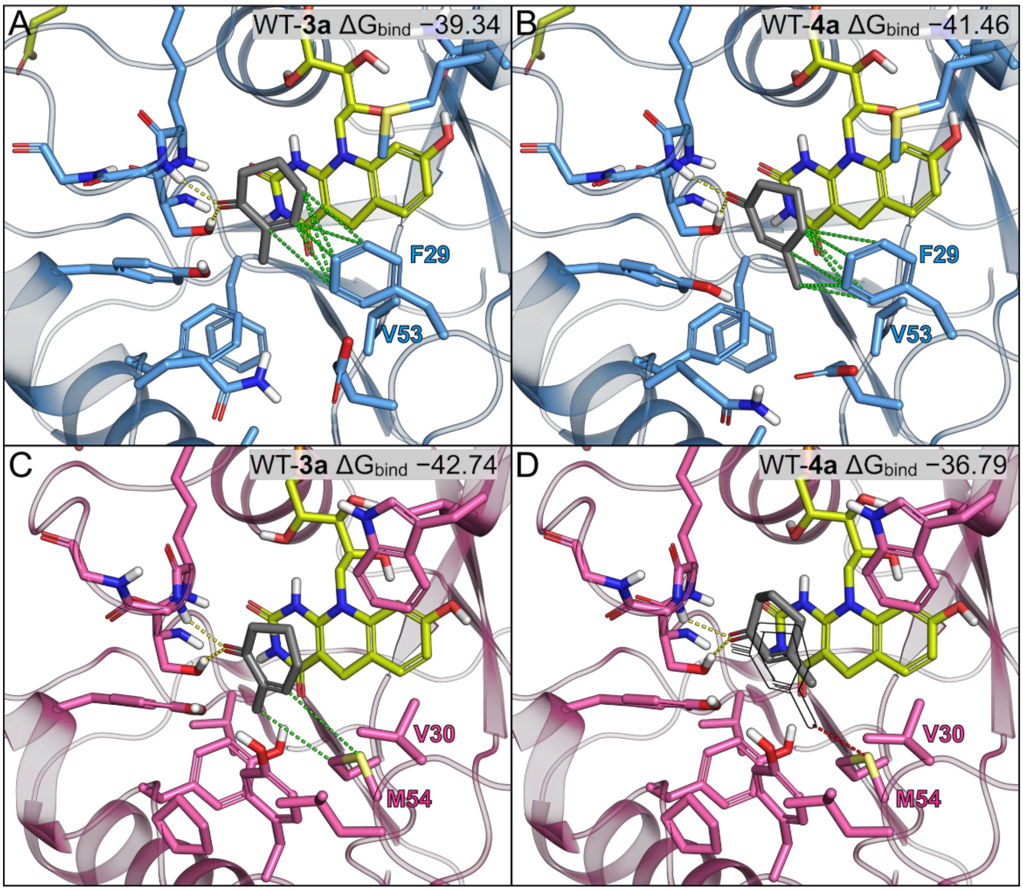

**Figure S13**. Docking poses of MSMEG_2027 and MSMEG_2850 with **3a** and **4a**

Induced fit docking poses were used to rationalize the observed stereochemistry with α- or β-substituted substrates. MSMEG_2850 and MSMEG_2027 are colored blue and pink, respectively. Hydrogen bonds are represented by yellow dashed lines, and relevant hydrophobic interactions by green dashed lines. A steric clash is indicated by a red dashed line with an imaginary favourable position of **4a** (black outline, D).

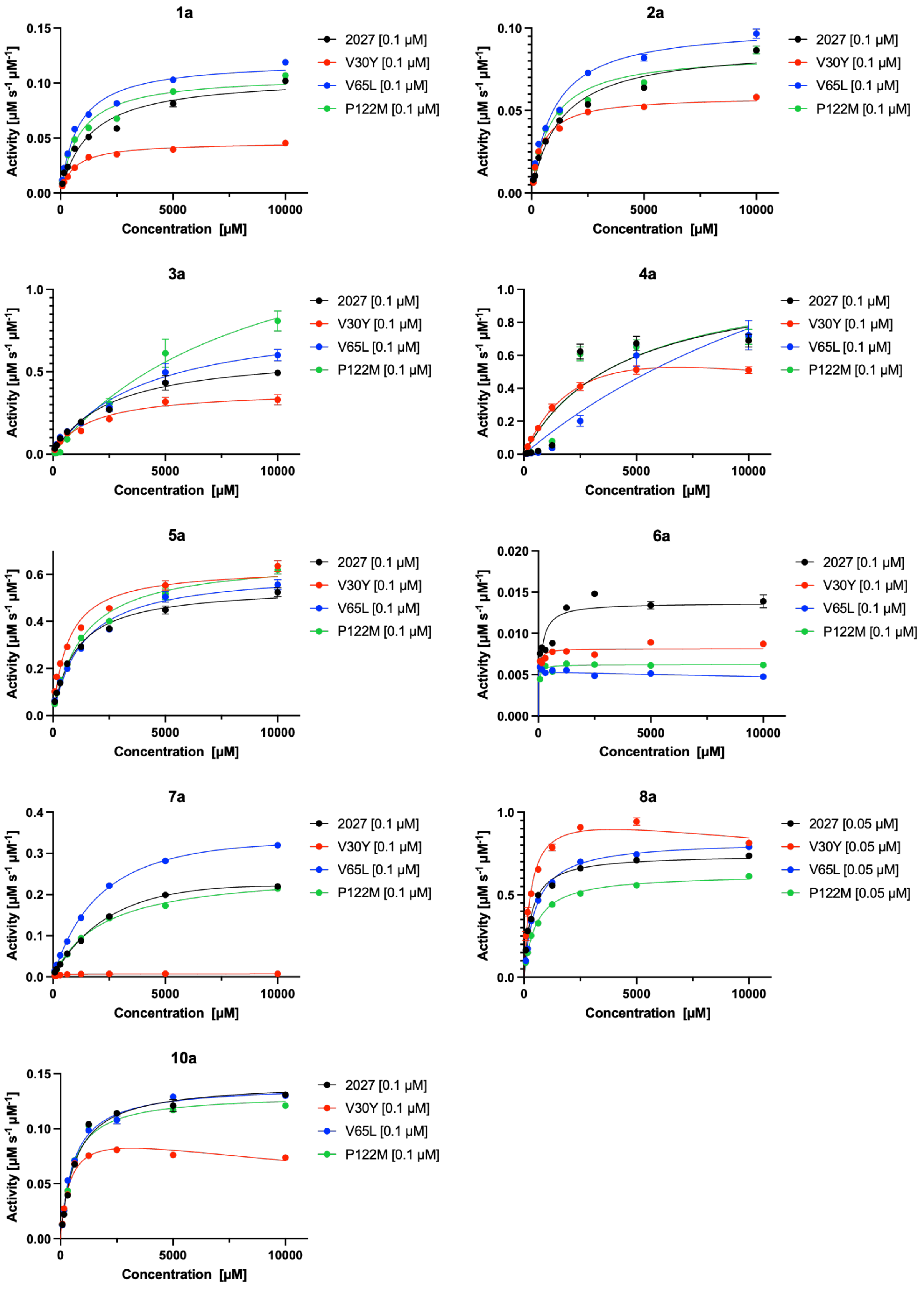

**Figure S14.** Steady state kinetics for MSMEG_2027 variants

Assays were performed using 50 mM tris buffer (pH 8.0), 10 mM $F_{420}$, 10,000–78.125 µM substrates, 0.05–0.1 µM enzyme.

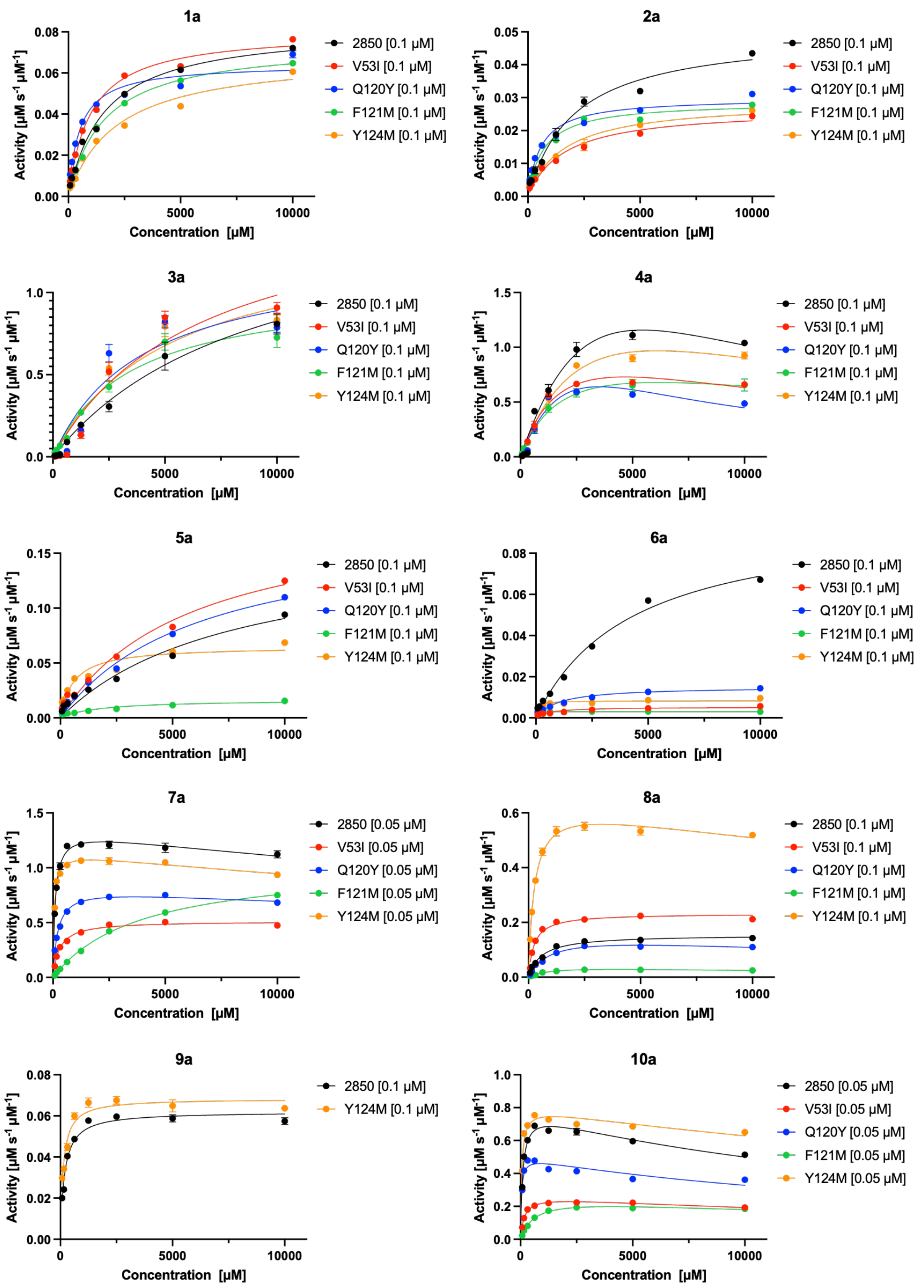

**Figure S15.** Steady state kinetics for MSMEG_2850 variants

Assays were performed using 50 mM tris buffer (pH 8.0), 10 mM $F_{420}$, 10,000–78.125 µM substrates, 0.05–0.1 µM enzyme

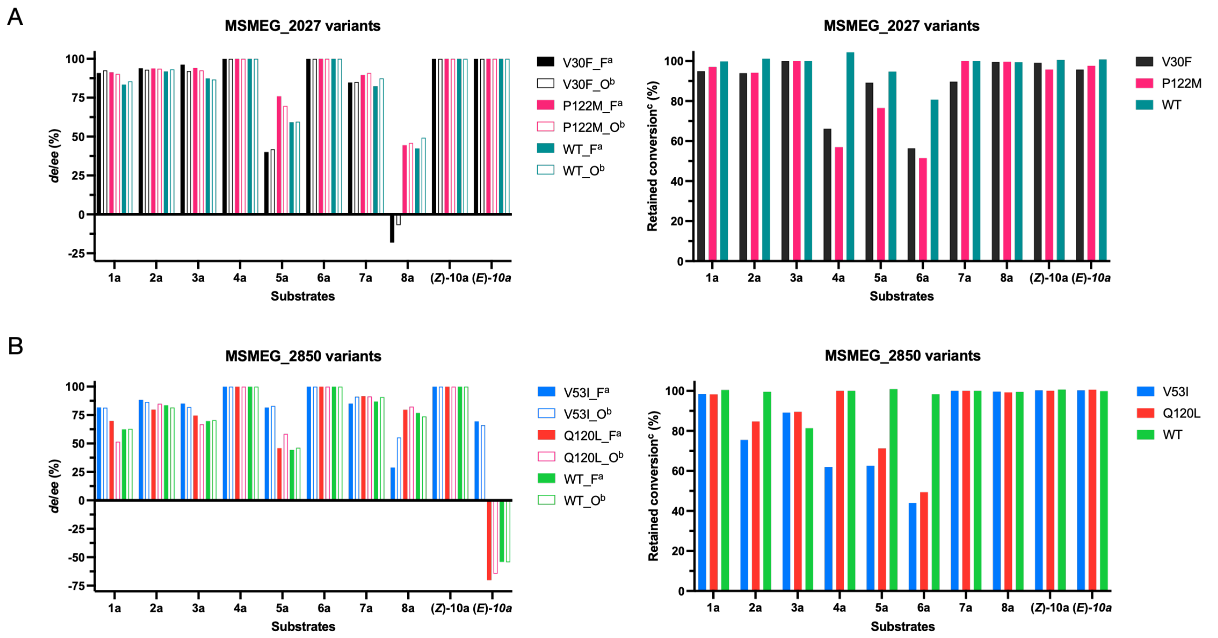

**Figure S16.** Residual activity and selectivity of variants following long terms storage at 4 °C Variants of MSMEG_2027 (A) and MSMEG_2850 (B) were assayed following three months of storage with each substrate and the relative conversion (% of activity with freshly prepared enzyme) and absolute stereoselectivity were determined.

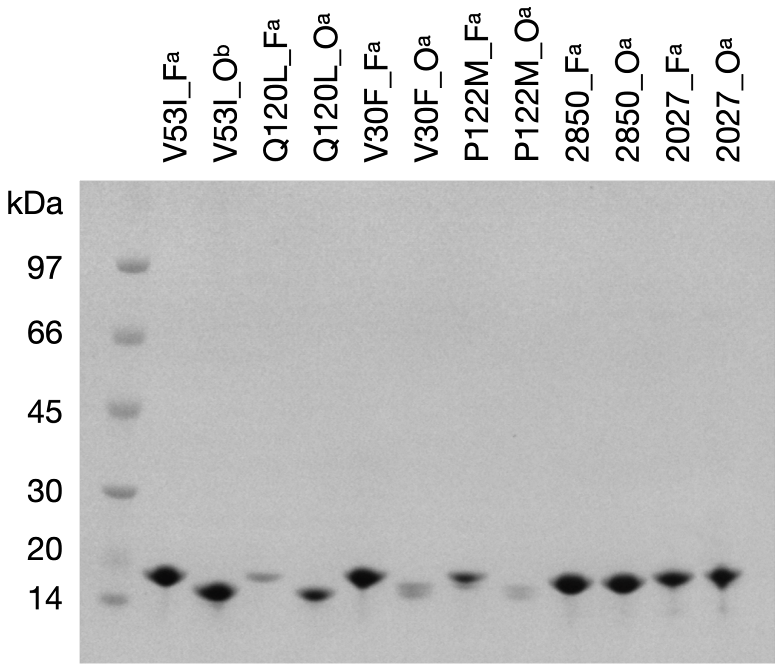

**Figure S17.** SDS–PAGE of enzymes freshly prepared and stored at 4°C for three months
The left lane shows the molecular weight (MW) ladder (Amersham low molecular weight (LMW) marker, GE Healthcare). [a] Freshly prepared enzyme. [b] Enzyme stored at 4°C for three months.
